# Corrective sub-movements link feedback to feedforward control in the cerebellum

**DOI:** 10.1101/2025.08.01.668191

**Authors:** Courtney I. Dobrott, Matthew I. Becker, Abigail L. Person

**Affiliations:** Neuroscience Graduate Program, University of Colorado Anschutz; University of Colorado Anschutz Department of Physiology and Biophysics; University of Washington Department of Psychiatry and Behavioral Sciences, Psychiatry Residency Research Program

**Keywords:** cerebellum, feedback control, feedforward control, error correction, forward model, corrective sub-movements, reaching, optimal feedback control, adaptation, motor learning

## Abstract

The ability to execute accurate movements is thought to rely on both anticipatory feedforward commands and rapid feedback corrections, yet how these control systems are integrated within cerebellar circuits remains unclear. Here, we show that corrective sub-movements (CSMs) – structured, feedback-driven adjustments occurring spontaneously during naturalistic mouse reaching – are not only encoded in anterior interposed (IntA) output neurons, but also act as instructive signals for feedforward learning. Closed-loop perturbations that trigger CSMs lead to learned shifts in the timing of future corrections, even in the absence of further perturbation. Strikingly, this learning depends on the timing of corrective responses, rather than the timing of the error itself, and is accompanied by physiological adaptation in cerebellar output neuronal firing rates. These findings reveal a cerebellar mechanism by which feedback responses train future anticipatory control signals, bridging the gap between reactive and anticipatory motor control in the cerebellum.

**Key Findings/ Highlights:**

- Mice generate rapid, precise corrective sub-movements (CSMs) during reach that counter both spontaneous variability and induced errors
- Cerebellar output neurons encode both predictive and corrective movements, mechanistically linking feedforward and feedback control
- Closed-loop optogenetic disruption of cerebellar output induced reach errors that were entirely compensated by CSMs
- Learning was driven by the timing of corrective responses, not the timing of the initial error, suggesting an instructive role for CSMs
- Both reach kinematics and cerebellar nuclear activity adapted over trials of repeated perturbation
- Optogenetic activation of cerebellar output drove errors and subsequent learning, but also blocked expression of that learning acutely, pinpointing cerebellar output as a key site of motor learning expression
- Findings suggest a neural mechanism by which feedback corrections influence learning of feedforward control policies in cerebellar circuits

## Introduction

A key feature of motor control is the ability to rapidly correct movements in response to errors ^1–3^. Motor cortex and spinal circuits are well-established mediators of rapid error correction, with spinal reflexes supporting fast, automatic responses and motor cortex contributing to longer-latency goal directed adjustments ^4–11^. In contrast, the cerebellum’s role in feedback control is less clearly delineated. Patients with cerebellar damage typically remain capable of using feedback to guide movements, selectively losing predictive, feedforward control ^12–17^. Nevertheless, the nature of feedback control in cerebellar patients is altered, showing phase lags, overcorrections, and an inability to adapt to feedback, that suggest interactions between feedforward and feedback control by the cerebellum ^17–21^.

Evidence for rapid error correction by cerebellum has also been seen in saccade perturbation experiments, in which induced errors are compensated by secondary saccades that accurately bring the eye back to target ^20,22–24^. Because of the speed of these corrective movements, it was argued that compensatory movements utilize internal feedback loops that rely on feedforward control signals (e.g. ‘efference copies’) in addition to sensory feedback ^20,22–24^. In this way feedback control may be linked with cerebellar feedforward control of movements.

Insight into how feedforward and feedback signals may be mechanistically linked comes from work identifying a region of cerebellar nuclei that contributes to anticipatory control of mouse reaching movements ^31–34^. Many neurons in the anterior interposed nucleus (IntA) burst near reach endpoint and exert an inward pull on the limb. Closed-loop inactivation of these neurons causes over-reach, while activation drives under-reach, revealing a causal role for reach-aligned neural activity ^31,32^. IntA neurons are under the control of Purkinje cells (PCs) which undergo adaptation in response to repeated perturbations of cerebellar inputs ^35^. Similarly, PCs adapt to novel forces applied to manipulanda in non-human primates (NHP), and show lead-and-lag relationships suggestive of feedforward and feedback control ^36–38^. These data raise the question of whether the physiological signals in cerebellar outputs represent feedback control – homing the limb to the target with fine online adjustments; feedforward control – using cerebellar inputs as cues to generate predictive commands^35,39^; or both.

Here we asked whether IntA output neurons encode corrective movements in addition to feedforward control signals. We found that mice exhibit corrective sub-movements (CSMs) during skilled reaching, indicating that real-time, feedback-based correction is a fundamental component of reach execution. These CSMs were functionally corrective, as they were consistently preceded by deviations from typical reach trajectories and contributed to maintaining endpoint accuracy. Recordings of IntA neurons during reaching revealed strong modulation during the reach, consistent with feedforward control, with additional modulation associated with CSMs. We then used closed-loop neural perturbations of red nucleus (RN) projecting IntA neurons (IntA^RN^) to perturb movements and found that mice generated fast, error-magnitude scaled CSMs. Interestingly, the timing of CSMs, not error, drove subsequent experience-dependent changes in CSM timing of unperturbed movements. Finally, we found that repeated IntA^RN^ excitation drove adaptation, both behaviorally and neurally, that was evident after stimulation was stopped, suggesting covert learning upstream of the cerebellar nuclei. Together, these findings link the generation of corrections to the learning of feedforward control signals, and show that these signals are superimposed in cerebellar output.

## Results

### Reaching movements in mice contain spontaneous CSMs

To enable mechanistic investigation of cerebellar contributions to CSMs, we first established whether and when discrete CSMs occur during unconstrained reaching in mice. Freely behaving mice were trained to perform a self-initiated skilled reaching task, retrieving a pellet from a pedestal, while 3D paw trajectories were recorded using high-speed cameras and motion capture from a reflective marker affixed to the wrist (***Fig. 1A***). As reported previously ^40,41^, time-averaged speed profiles exhibited a characteristic asymmetric bell-shaped curve with a slightly extended decelerative phase compared to the accelerative phase (***Fig. 1B***). Because trial averaging can obscure variably timed corrections, we analyzed single-trial kinematics for the presence and timing of CSMs relative to a mid-reach positional landmark termed “threshold” (***Fig. 1C-D***). During reach acceleration, CSMs were identified as peaks in jerk (the third derivative of position), indicating transient increases in positive acceleration. During reach deceleration, CSMs were defined as periods of positive acceleration lasting more than 25 ms within a 200 ms window following threshold crossing (***Fig. 1D, Supplemental Fig. 1***).

**Figure 1:**
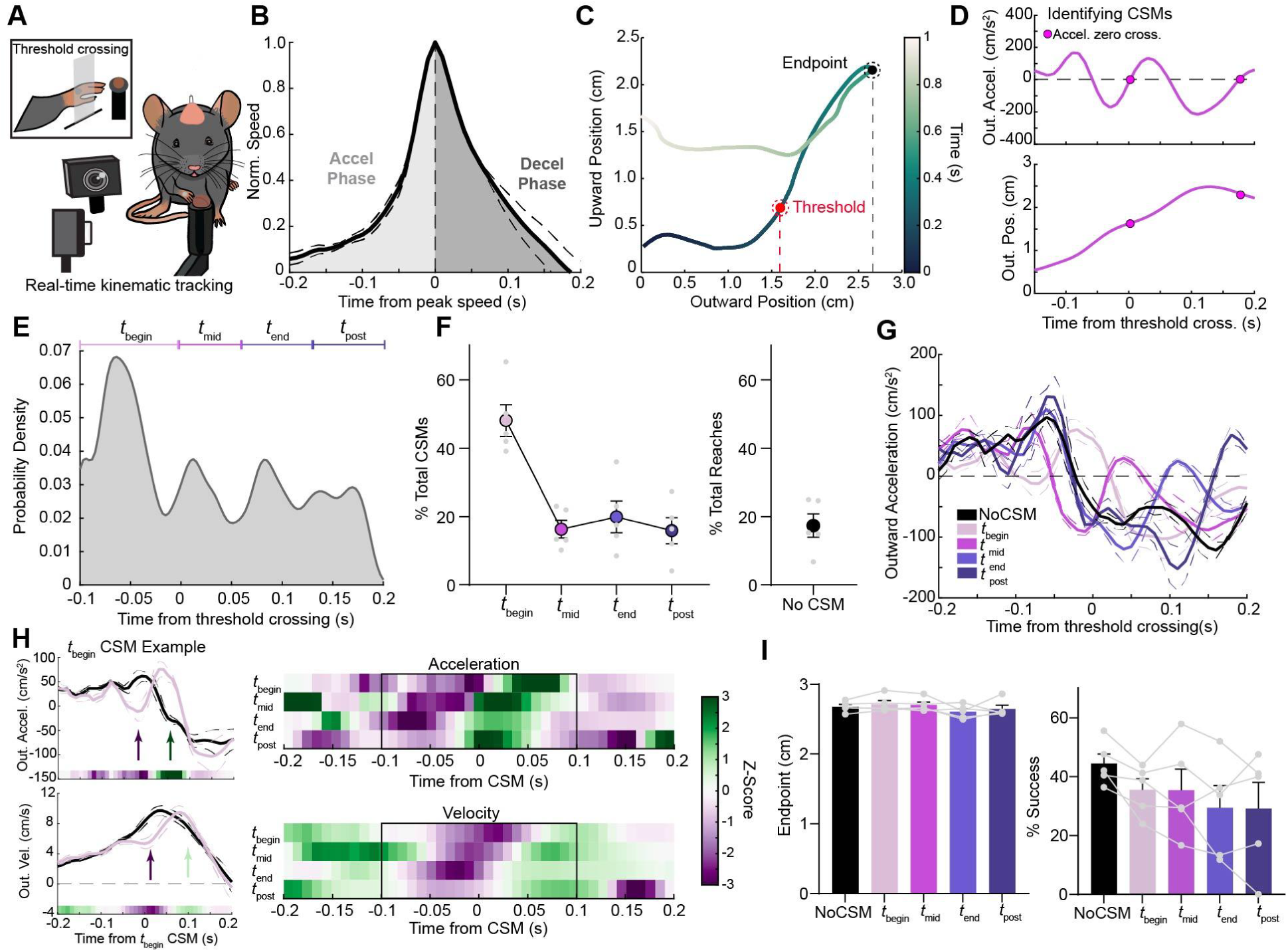
Reaching movements in mice exhibit spontaneous and structured CSMs. (A) Schematic of behavioral setup for freely behaving reaching. (B) Normalized speed profile for control (Sham) mice (N=5 mice, n=2074 reaches). Average reach speed aligned to peak. Thin dashed lines: ±SE ; bold lines: mean across mice. (C) Example single reach position trajectory illustrating threshold crossing (red dashed line) and final endpoint (black dashed line). Colorbar represents time. (D) Example outward position (top) and acceleration (bottom) trajectory aligned to threshold crossing. Pink dots identify zero-crossings in acceleration at CSM onset. (E) Probability density function of CSM timing distributions for all mice aligned to threshold crossing. (F) Left: Summary data showing percentage of CSMs occurring in each epoch. Right: Percentage of reaches with no CSMs. Individual mice, small dots; averages (±SEM), large dots. (G) Average acceleration for reaches with no CSMs (black) or reaches with CSMs that fell within each epoch (light purple to dark purple), aligned to threshold crossing. Thin dashed lines: ±SE ; bold lines: mean across mice. (H) Left: Example kinematics showing average acceleration (top) and velocity (bottom) trajectories of averaged t_begin_ CSM-containing reaches (pink), aligned to t_begin_ CSM onset or all reaches aligned to t_begin_ CSM (black; template). Arrows indicate deviations from mean trajectories and corrections. Heatmaps below trace same as the Right panel. Right: Pointwise z-score heat-map comparing acceleration (top) or velocity (bottom) of template reaches versus CSM-containing reaches, for each epoch. Note that t_begin_ CSMs are defined by jerk rather than acceleration, accounting for the small rightward shift in green (CSM) from 0. (I) Left: Average endpoint position for reaches with or without CSMs separated by CSM time (RM One-Way ANOVA, *p* = 0.18, *N* = 5 mice). Right: Percentage of successful reaches across groups (RM One-Way ANOVA, *p* = 0.11, *N* = 5 mice). Light gray circles and lines represent individual mice; bars, group mean ± SEM.

When kinematics were examined on a reach-by-reach basis, CSMs were common, with only 17% of all reaches lacking them (***Fig. 1E-F***, right). Interestingly, the timing of CSMs was nonuniform, showing regular peaks in probability that occurred approximately every 70 ms, reminiscent of periodic CSM timing distributions seen in human reaches (***Fig. 1E***) ^42,43^. For further analysis, we leveraged this natural multimodal distribution to classify CSM times into epochs spanning the beginning, middle, and end of outreach as well as return, termed t_begin_, t_mid_, t_end_, and t_post_, respectively. Among reaches containing CSMs, nearly half (48%) exhibited CSMs during the initial acceleration phase (t_begin_; ***Fig. 1F,*** left), with the remaining reaches containing CSMs occurring during later movement phases. These phase-specific CSMs can be visualized in averaged kinematics of reaches grouped by CSM timing, aligned to the positional threshold crossing (***Fig. 1G, Supplemental Figure 1***).

Although these accelerative bouts resembled CSMs described for other movements ^4,5^, we wanted to determine whether they were truly corrective. We reasoned that if they were corrective, the kinematics preceding the CSM should deviate from the average trajectory, indicating an error to correct. To test this, we grouped reaches by CSM timing (e.g. t_begin_, etc) and compared CSM-aligned kinematics to a ‘template’ that was the average and standard deviation of all reaches aligned to the reference epoch, using a sliding-window z-score analysis (***Fig. 1H,*** *Methods*). As expected, z-scores were high during the CSM itself (green in the acceleration heat maps), reflecting the higher-than-average acceleration. Notably, we also observed negative z-score deviations in kinematics before the CSM (purple), indicating a deviation from the average trajectory (***Fig. 1H***, top, and arrows). These divergences from the mean both preceded CSMs and were in the opposite direction of the CSMs, suggesting that CSMs act to correct ongoing errors and restore the trajectory toward the mean. Indeed, despite these kinematic variations, velocity moved toward the mean following CSMs (***Fig. 1H,*** bottom). While reaches that contained CSMs were associated with a moderate decrease in success rate overall (paired t-test, *p=0.048*), success rates did not differ significantly across each CSM epoch (***Fig. 1I*** ; RM One-way ANOVA, *p=0.11*, N=5 animals). Together, these findings suggest that CSMs compensate for online deviations in reach kinematics, supporting accurate targeting.

### IntA neurons encode both feedforward reach control and CSMs

IntA neuron firing rates scale limb kinematics during reaching, contributing to reach precision and accuracy ^32^. However, it remains unclear whether IntA activity reflects corrective movements that steer the limb to the target or functions primarily as a feedforward, anticipatory brake. To evaluate this question, we first established that CSMs occurred in reaches from head-fixed mice, a configuration that facilitates acute in vivo-recordings (***Fig. 2A***). As in the freely behaving condition, reaches made in the head-fixed configuration also contained CSMs with a non-uniform distribution, although with just two prominent peaks, such that the majority of CSMs occurred during the t_begin_ (52%) and t_end_ (33%) epochs (***Fig. 2B***; ***Supplemental Fig 2A-E***).

**Figure 2:**
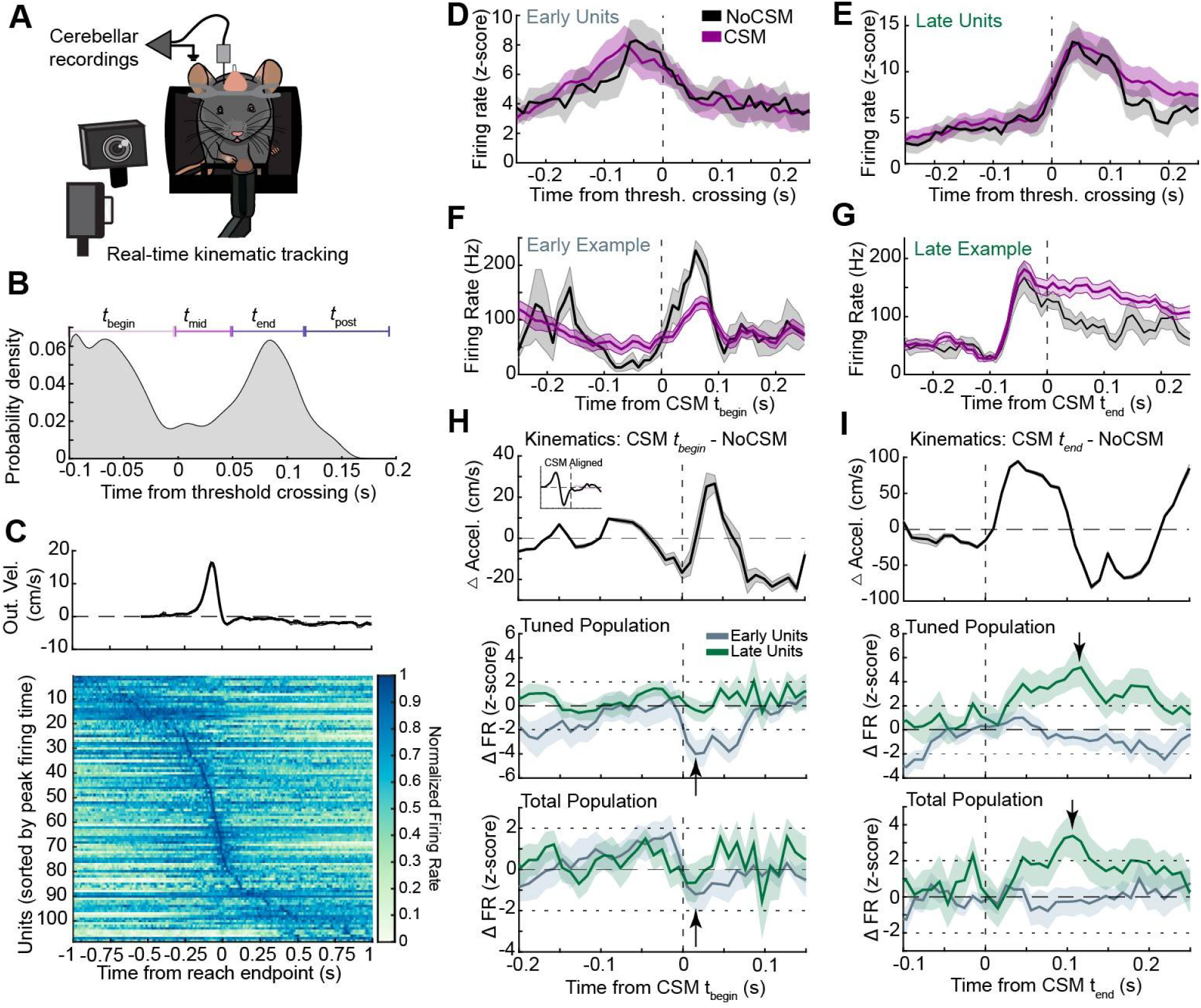
IntA neurons encode both feedforward reach control and CSMs. (A) Schematic of Neuropixel recordings in the IntA during head-fixed reaching. (B) Probability density function summarizing distribution of CSM times in head-fixed reaches (n = 1558 reaches, N=10 mice), showing multimodal structure. (C) Top: Average outward velocity trace aligned to reach endpoint across all trials. Bottom: Heatmap of normalized firing rates for all recorded IntA neurons, sorted by time of peak activity, aligned to reach endpoint (n=108 units, N=6 mice) (D-E) Z-scored population firing rates across all significantly modulated units (Early Units: n=23 neurons, Late Units: n=21 neurons, N=6 mice) (F-G) PSTHs from example neurons aligned to CSM t_begin_ onset (Early Unit Example) and CSM t_end_ onset (Late Unit Example). (H) **Top:** Average difference in acceleration (CSM t_begin_ − NoCSM) aligned to CSM_begin_ onset. Inset shows schematic of alignment strategy. **Middle:** Z-scored population firing rate differences between reaches with and without CSMs. Early (blue) and late units (green) with CSM specific modulation aligned to CSM_begin_ onset (Early Units, n=11 out of 23 neurons; Late Units, n=15 out of 21 neurons; Early units arrow, z= −4, p<0.0001 (two-tailed z-test)). Thin line is the mean and shaded regions indicate SEM. Small horizontal dashed lines indicate ±2 z-scores, corresponding to approximately the 95% confidence interval under a standard normal distribution. **Bottom**: Z-scored population firing rate difference for all Early and Late Units (Early units, n=23 neurons; Late units, n=21 neurons; Early Units arrow, z= −1.19, p=0.23 (two-tailed z-test). (I) **Top**: Same as H but aligned to CSM t_end_ onset. **Middle**: Z-scored population firing rate differences between reaches with and without CSMs. Early (blue) and late units (green) with CSM specific modulation aligned to CSM_end_ onset (Early Units, n=11 out of 23 neurons; Late Units, n=15 out of 21 neurons; Late Units arrow, z= 5.19, p= 2×10^-7^ (two-tailed z-test). **Bottom**: Z-scored population firing rate difference for all Early and Late units (Early units, n=23 neurons; Late units, n=21 neurons; Late Units arrow, z= 3.38, p=0.0007 (two-tailed z-test).

To address whether neurons were differentially modulated during CSMs, we performed acute Neuropixel recordings of IntA during head-fixed reaching. In stereotaxically targeted recordings, units were included for further analysis if they were located within the expected depth range of the IntA, confirmed by posthoc histological analysis, and showed significant firing rate changes during reach. Consistent with previous findings ^32^, many IntA neurons exhibited robust modulation near reach endpoint (***Fig. 2C***, n=108 units, N=6 mice***, Supplemental Fig 2F***). We segregated neurons for subsequent analyses based on whether their trial-averaged peak firing rate occurred within a 100 ms window before (Early Units) or after peak outward velocity (Late Units), accounting for the primary peaks in the distribution of CSM times.

To test for neural correlates of CSMs, we compared neural activity during reaches with and without CSMs. Notably, at the population level for both Early and Late units, IntA reach-related modulation was largely the same between CSM-containing and No-CSM reaches (***Fig. 2D-E***), suggesting that the primary signal from cerebellar output neurons is not related to CSMs. However, aligning to t_begin_ or t_end_ epochs, we found subsets of neurons that were differentially modulated during CSM-containing reaches (***Fig. 2F-G***). Among Early Units aligned to t_begin_ CSM onset, 11 out of 23 neurons showed significantly reduced firing rates during reaches containing a t_begin_ CSM compared to reaches with no CSM. In this subpopulation, firing rates dropped an average of 25.3 ± 6.6 spikes/s below rates during No-CSM reaches, with a maximum difference 15 ms after CSM onset (***Fig. 2H,*** middle, *arrow,* z = −4, *p <0.0001* (two-tailed z-test)). Relatedly, among Late Units aligned to t_end_ CSMs, 15 out of 21 neurons fired significantly stronger during t_end_ CSM-containing reaches relative to No-CSM reaches. These neurons showed an average increase of 23.8 ± 6.6 spikes/s over No-CSM reaches, peaking 116 ms after CSM onset (***Fig. 2I,*** middle, *arrow,* z = 5.2, *p =2.1×10^-7^*), as the CSM itself decelerated. Interestingly, neither Early nor Late populations showed significant rate modulations during CSMs in the converse epoch alignment (***Fig. 2H-I,*** middle, gray vs green traces).

These observations were only partially robust at a total population level: Averaging across all Early Units, firing rates dropped slightly, but non-significantly, relative to No-CSM reaches as t_begin_ CSMs started, averaging −9.6 ± 4.7 spikes, peaking 15 ms after CSM onset (***Fig. 2H,*** bottom, *arrow,* z = −1.2, *p = 0.2*) and were only weakly modulated around t_end_ CSMs. In contrast the firing rate changes observed in Late Units during CSMs were robust at the population level. Late Units as a whole exhibited significant increases in activity following t_end_ CSMs (***Fig. 2I,*** bottom, *arrow,* z=3.38, *p= 0.0007*), averaging 13.0 ± 6.8 spikes, peaking 106 ms after CSM. Given previous work implicating IntA activity in anticipatory control of limb deceleration ^32^, these data are consistent with the hypothesis that both Early and Late Units operate along the same control axis (inward pull on the limb), with Early Units associated with the accelerative component of the CSM, potentially through a reduction of inward pull, followed by an increase in decelerative pull exerted by the Late units. Together, these findings show that IntA neurons primarily signal predictive decelerative commands, with superimposed, small signals encoding corrective movements.

### CSMs compensate for induced errors during goal-directed reaches

The observations above suggest that cerebellar output contains both anticipatory and feedback control signals. However, if the cerebellum itself induces error, can the brain self-correct within the same reach? We tested this idea directly with closed-loop optogenetic perturbations of IntA premotor output neurons that project to the RN, which comprise the primary premotor output population from IntA and collateralize to the thalamus and other brainstem structures. We introduced ChR2 into IntA neurons that project to the RN (IntA^RN^) using an intersectional genetics approach in which Vglut2-cre animals were injected with AAV-Con/Fon-hChR2 in the IntA and AAVretro-FlpO in the contralateral RN (***Fig. 3A-B***), and an optical fiber targeting ipsilateral IntA ^44^. We then performed closed-loop stimulation of these neurons during reaching, triggering a 50 ms train (2 ms duty cycle; 100 Hz) at a prespecified positional threshold on a randomized 25% of trials.

**Figure 3:**
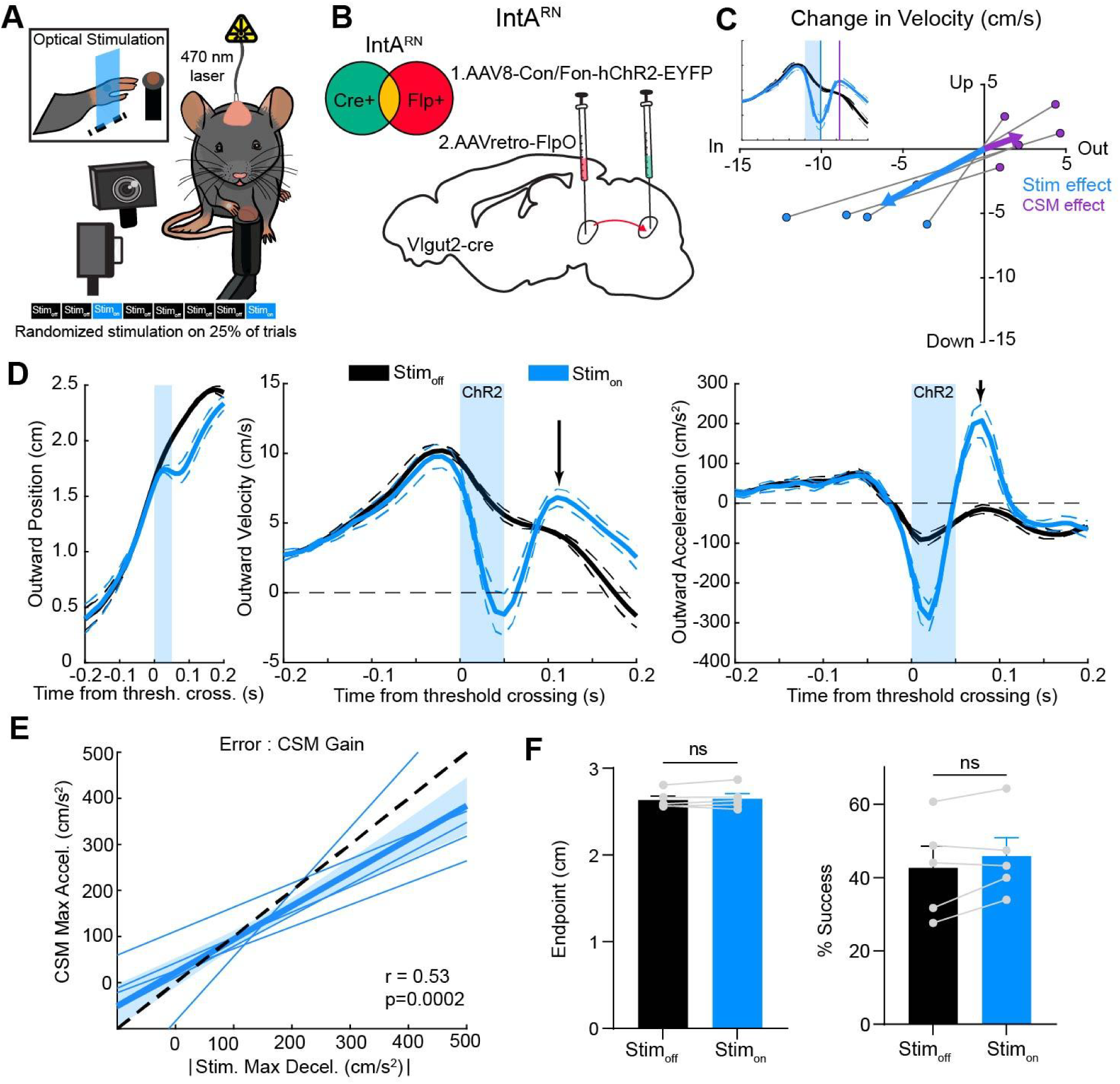
CSMs compensate for induced errors during goal-directed reaches. (A) Schematic of behavioral setup with real-time kinematic tracking and closed loop optogenetic stimulation. Stimulation at positional threshold (1.6 cm) on a randomized 25% of trials, 470 nm laser with a stimulation train duration of 50 ms (2 ms pulses, 100 Hz). Stim_on_, (laser-on) reaches, Stim_off_, (laser-off) reaches. (B) Intersectional genetics approach for targeting excitatory IntA neurons that project to the RN. (C) Average change in outward and upward velocity for each animal at (Stim_on_ - Stim_off_) 50ms after threshold crossing (Stim effect) and 110 ms after threshold crossing (CSM effect). The average effect size is indicated by arrow length (Stim effect, *p=0.003* (outward, −7.09 ± 1.56 cm/s, upward, −4.87 ± 0.54 cm/s), CSM effect, *p=0.01*, one sample t test (outward, 2.67 ± 0.78 cm/s, upward 1.22 ± 0.85 cm/s)). Comparison between groups (Outward: *p=0.0005*, Upward *p=0.0003*, two sample t-test). Gray lines show matched data per animal. (D) Kinematic profiles of outward position (left), velocity (middle), and acceleration (right) aligned to threshold crossing (Stim, blue shaded region). Traces compare trials with no stimulation (Stim_off_, black) and stimulation (Stim_on_, blue), arrows mark the CSMs. Data represent mean ± SE across N=5 mice. (E) Across mice, greater optogenetically induced deceleration (x-axis: max outward deceleration during stimulation epoch) was strongly associated with compensatory CSM responses (y-axis: max outward acceleration during CSM) (r = 0.53, p=0.0002, Fisher z t-test, N=5 mice). Individual lines represent the mean of each animal, and the thick line represents the group mean ± SEM across mice. The dashed line indicates unity. (F) Endpoint position (left) (p>0.05). Success rate (right) (p > 0.05, paired t-test), and in 3 of 5 mice it increased in stimulated trials. Gray lines show matched data per animal. N=5 mice.

Closed loop optogenetic stimulation of IntA^RN^ neurons during reach induced a rapid decrease in outward and upward velocity (Stim effect: outward, −7.1 ± 1.6 cm/s, upward, −4.9 ± 0.5 cm/s), mirroring the effects observed with pan-neuronal IntA (IntA^All^) stimulation in previous work ^32^, which we reanalyze here for purposes of comparison (***Fig. 3C, Supplemental Fig 3B;*** Stim effect: outward, −6.3 ± 1.3 cm/s, upward, −7.0 ± 1.1 cm/s; See Methods). Acute effects during optogenetic stimulation were statistically indistinguishable between the IntA^RN^ and IntA^All^ stimulus conditions (Stim effect: outward velocity change *p=0.72*, upward velocity change *p=0.18*, two-sample t-test). However, following stimulation, a clear difference emerged across these conditions: Following IntA^RN^ stimulation-induced deceleration, mice consistently compensated with a rapid and large reacceleration (***Fig. 3C-D,*** arrows, CSM effect (avg. time of max outward velocity for the CSM (110 ms)): outward velocity, 2.7 ± 0.8 cm/s, upward velocity, 1.2 ± 0.9 cm/s) that redirected the limb back to the target, which was inconsistent in the IntA^All^ stimulus condition (***Supplemental Fig. 3C;*** CSM effect: change in outward velocity: −2.0 ± 1.4 cm/s, upward velocity, −0.9 ± 0.9 cm/s).

Given the speed and consistency of the induced CSMs following IntA^RN^ stimulation, we next asked whether these induced CSMs scaled with the magnitude of the induced error, which would suggest online monitoring of reach displacement. To test this, we examined the relationship between stimulation induced deceleration and the resulting CSM acceleration. We found a strong positive correlation between acute stimulation-induced deceleration magnitude and CSM reacceleration magnitude (***Fig. 3E,*** r=0.53, *p=0.0002*). We also found that reach endpoints and success rates were unchanged following IntA^RN^ stimulation compared to unstimulated reaches (***Fig. 3F,*** Endpoint (left): p>0.05, SuccessRate (right): p>0.05, paired t-test), indicating CSMs effectively scale with and compensate for acute perturbations. These findings differed from pan-neuronal stimulation IntA^All^ reported previously, in which animals consistently undershot the target following stimulation ^32^. To examine the difference between corrective responses across stimulus conditions further, we analyzed the timing and scaling of CSMs following IntA^All^ stimulation. CSM reacceleration scaling was much weaker relative to the acute deceleration from stimulation (***Supplemental Fig. 3E,*** *r=0.34, p=0.009;* IntA^All^ vs IntA^RN^, p<0.0001, two-sample t-test). The lower gain of these CSMs was accompanied by significantly more hypometric endpoints (***Supplemental Fig. 3F,*** Endpoint (left): *p=0.004*, Success Rate (right): *p=0.1*, paired t-test). In summary, CSMs following IntA^RN^, but not IntA^All^, stimulation facilitated precise and accurate endpoints through fast, scaled corrections, suggesting that the mouse motor system may perform computations that estimate limb displacement within a given reach to support fast corrections.

### CSM response timing, not error timing drives learning

Optimal feedback control frameworks propose that rapid feedback corrections are used to update future movements through learned predictive strategies ^7,27,45,46^. We therefore asked whether induced CSMs influenced anticipatory control. Because unperturbed reaches in naive mice show distributed CSM timing, while induced CSMs were pegged to stimulus timing, we tested whether exposure to timed errors shifted the natural distribution of CSM timing. We compared trials in which we induced CSMs (Stim_on_) versus interspersed trials in which we did not stimulate (Stim_off_), using IntA^RN^ optogenetic activation. As expected, the distribution of CSMs in Stim_on_ reaches showed a stereotyped peak in CSM probability after the time of stimulation, with 51% of all CSMs occurring during the t_mid_ epoch and 21% during the t_end_ epoch (***Fig. 4A*** (middle) spanning latencies of 0 to 130 ms relative to threshold crossing). Interestingly, we observed a striking shift in the probability distribution of CSM timing for subsequent Stim_off_ reaches relative to naive conditions (Sham). In Stim_off_ trials, there was a doubling of the likelihood of CSMs occurring during the t_end_ epoch, increasing to 39% from 20% in the Sham condition (***Supplemental Fig. 4A,*** CSM timing (left) p<0.001, Kolmogorov–Smirnov test, Total CSM-containing reaches (middle) p=0.004, two-way ANOVA, multiple comparisons: t_begin_ p=0.03, t_end_ p=0.003).

**Figure 4:**
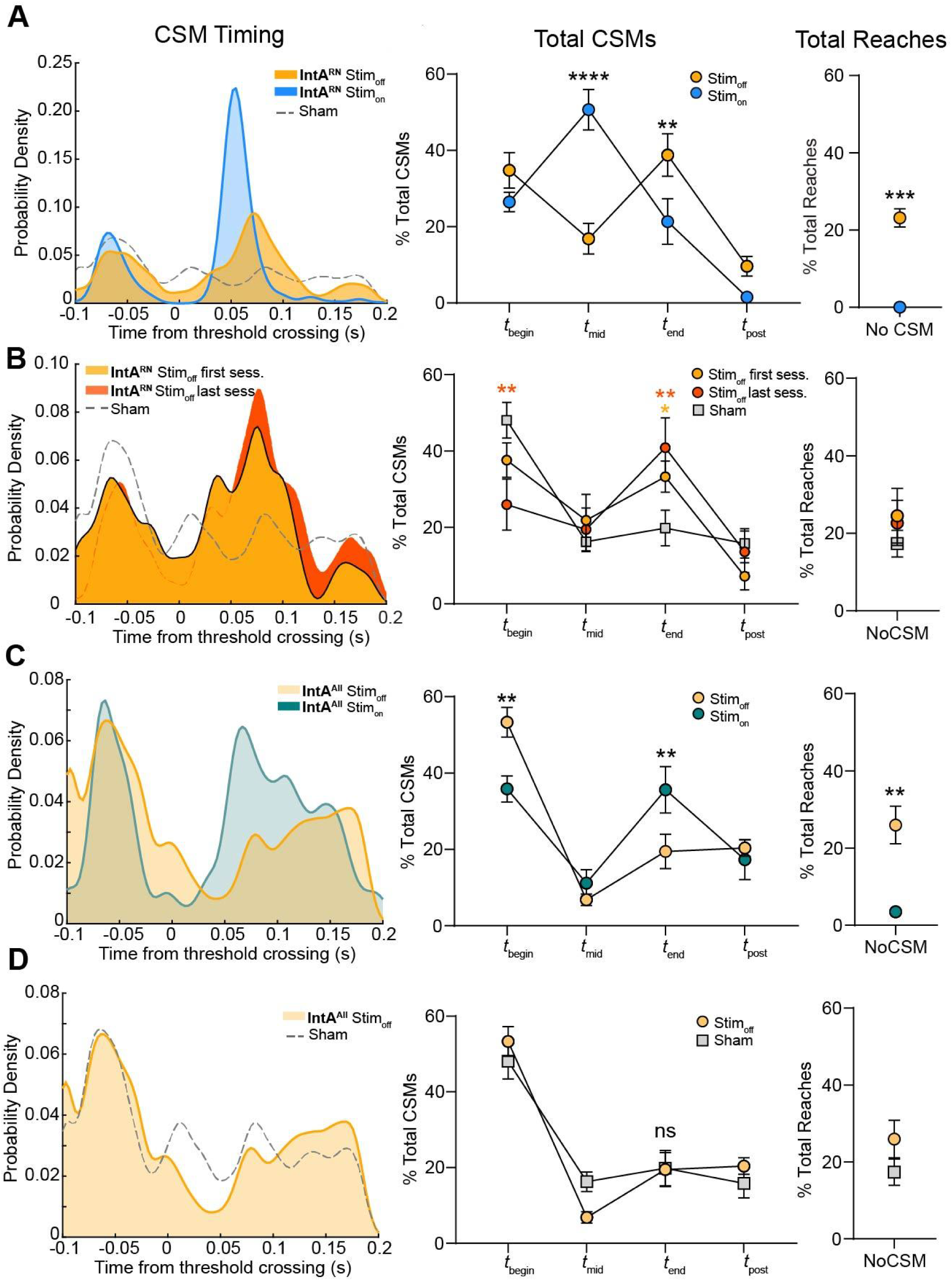
CSM response timing, not error timing drives learning. (A) ***Left***, Kernel density function of CSM timing for IntA^RN^ Stim_on_(blue) and interspersed Stim_off_ (yellow) reaches; Sham, dashed gray line. ***Middle***, Percentage of CSMs that occur within different epochs for IntA^RN^ Stim_on_ or Stim_off_ reaches. ***Right***, Percentage of reaches that had no CSM. CSM timing (p<0.001, D=0.35, Kolmogorov–Smirnov test), Total CSMs (p<0.0001, two-way ANOVA, multiple comparisons: t_mid_ p<0.0001, t_end_ p=0.007), Total Reaches containing CSMs (p=0.0006, paired t-test), N=5 mice. (B) Same as A but comparing the first (yellow) versus last session (orange) for IntA^RN^. CSM timing (first vs. last, p=0.09, D=0.14; first vs. sham, p=0.006, D=0.13, last vs. sham, p=0.0002, D=0.19, Kolmogorov–Smirnov test), Total CSMs (first vs. last, p=0.32; first vs sham, p=0.03, multiple comparisons: t_end_ p=0.04; last vs sham, p=0.004, two-way ANOVA, multiple comparisons: t_begin_ p=0.009, t_end_ p=0.007), Total Reaches containing CSMs (p>0.05, paired t-test). IntA^RN^ N=5 mice; Sham N= 5 mice. (C) Same as A but comparing IntA^All^ Stim_on_ (teal) and interspersed Stim_off_ (yellow) reaches. CSM timing (p<0.001, D=0.22,Kolmogorov–Smirnov test), Total CSMs (p=0.001, two-way ANOVA, multiple comparisons: t_begin_ p=0.003, t_end_ p=0.007), Total Reaches containing CSMs (p=0.004, paired t-test). N=8 mice. (D) Same as A but comparing IntA^All^ Stim_off_ (yellow) reaches and Sham control reaches. CSM timing (p=0.0006, D=0.08,Kolmogorov–Smirnov test). Total CSMs (p=0.18, two-way ANOVA). Total Reaches containing CSMs (p=0.2, paired t-test). IntA^ALL^ N=8 mice, Sham N=5 mice.

Notably, the shift in probability of CSM timing with exposure to induced errors emerged rapidly, appearing within the first session, where the probability of t_end_ epoch CSMs increased to 33% in Stim_off_ reaches, a significant increase from 20% in Sham controls (***Fig. 4B*** (middle) p=0.04, two-way ANOVA). Over-session accumulation of CSM timing consolidation was also evident: in the final session (6-11 sessions) the probability of CSMs occurring during the t_end_ epoch rose to 41% (p=0.007), whereas CSMs in the t_begin_ epoch dropped to 26% from 48% in Sham controls (***Fig. 4B*** (middle) p=0.009). These experience-dependent changes in CSM timing suggest that feedback responses influence feedforward control of subsequent reaches.

Is the shift in CSM timing distribution following IntA^RN^ stimulation driven by error timing, or CSM timing? To address this, we leveraged the fact that CSMs following IntA^All^ stimulation were slower and more temporally variable relative to those evoked by IntA^RN^ stimulation, while the induced errors had comparable magnitude and timing across stimulus conditions. CSM timing following IntA^All^ stimulation was less consolidated within the t_mid_ to t_end_ epochs, with only 47% of Stim_on_ reaches containing a CSM during this period, compared to 72% of Stim_on_ reaches in the IntA^RN^ condition (***Fig. 4 A,C***). Notably, the distribution of CSM timing in Stim_off_ reaches in the IntA^All^ condition did not show a significant shift compared to Sham controls (***Fig. 4D*** (middle) p=0.18, two-way ANOVA), despite the similar *error* timing and magnitude between IntA^All^ and IntA^RN^ stimulation. These findings suggest that the timing of the correction, not the error, drives learned CSM timing. This observation is conceptually similar to observations from human psychophysics, where correction timing influences learning ^46^.

### Neural correlates of induced CSMs

Having found that IntA neurons encode both anticipatory and spontaneous CSMs during reach (***Fig. 2***), we next asked whether the same neurons also encoded induced CSMs. To test this, we performed acute Neuropixels recordings during closed-loop stimulation of the IntA-to-RN (IntA^RN^) pathway. Stimulation was delivered distal to the recording site via an optical fiber targeting RN, following injection of AAV2-DIO-ChR2 into the IntA of Vglut2-Cre mice (***Fig. 5A-B***). This projection-specific approach allowed us to induce movement errors and corrective responses (***Fig. 5C,D,*** top) while minimizing optoelectric artifacts in recordings (***Fig. 5B***). Across mice, stimulation reliably altered reach kinematics: Stim_on_ trials exhibited a transient reduction in outward velocity during the stimulation period, followed by a stereotyped, induced CSM shortly after light offset (***Fig. 5C***, top; ***Fig. 5D***, top).

**Figure 5:**
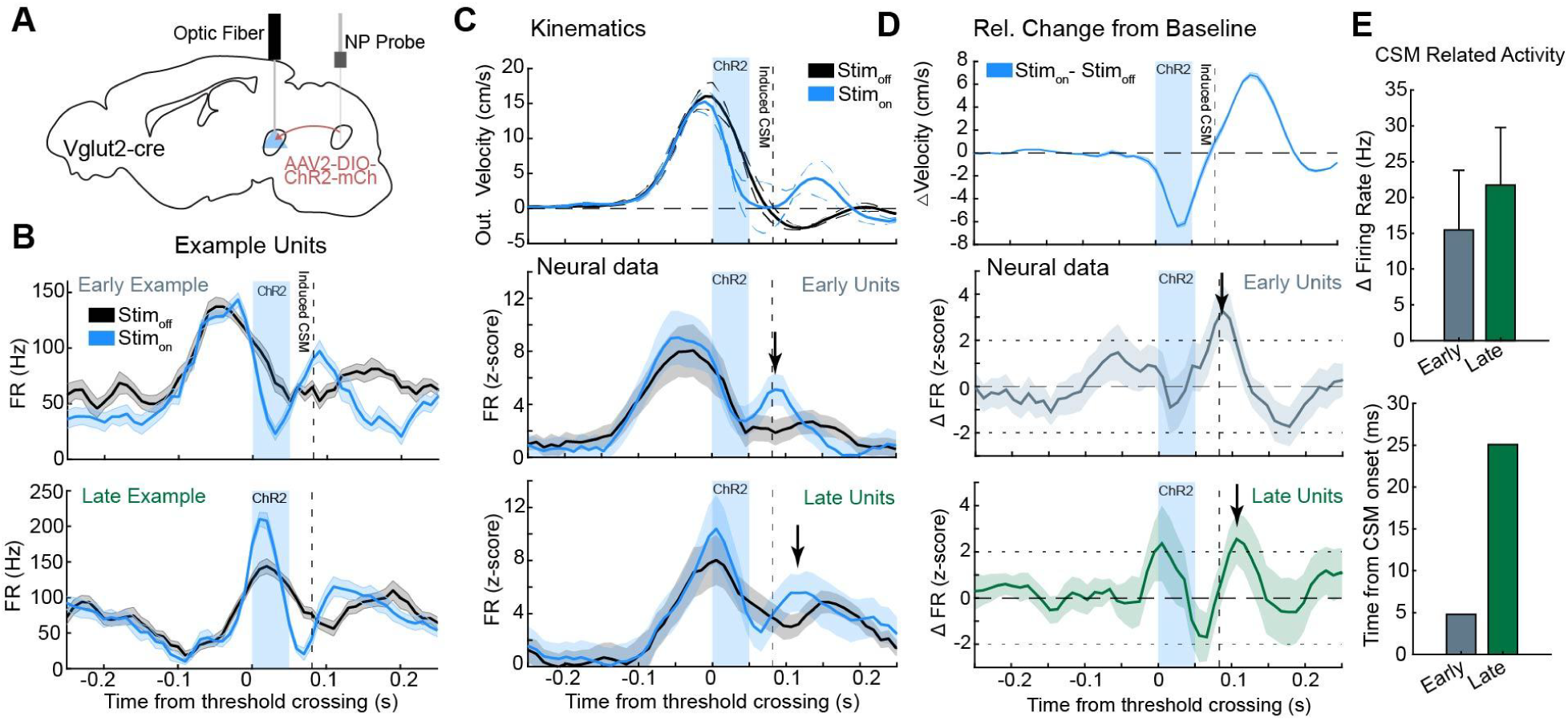
Neural correlates of induced CSMs. (A) Schematic of recording and optogenetic stimulation configuration during head-fixed reaching. (B) Firing rates from representative Early (**top**) and Late (**bottom**) Units during Stim_off_ (Black) versus Stim_on_ (Blue) trials aligned to threshold crossing. Dashed vertical lines indicate the average timing of the induced CSM (81.5 ms), N=3 mice. (C) Summary data showing outward velocity (**top**), and population average z-scored firing rates for Early Units (**middle**, n= 26 neurons) and Late Units (**bottom**, n=18 neurons). Shaded areas represent SEM and blue rectangle represents stimulation epoch. Arrows indicate significant changes in firing rate from baseline. (D) Relative change from baseline in velocity (**top**) and population average z-scored firing rate (middle, bottom) for Stim_on_ (blue) conditions, aligned to threshold crossing. **Middle**: Relative change in z-scored firing rate for Early Units (n=26 neurons, arrow, z= 3.26, *p=0.0012*). **Bottom**: Relative change in z-scored firing rate for Larly Units (n= 18 neurons, arrow, z= 2.56, p=0.01 (two-tailed z-test)). Thin lines show mean change ± SEM. Small horizontal dashed lines indicate ±2 z-scores, corresponding to approximately the 95% confidence interval under a standard normal distribution. (E) (**top**) Maximum change in firing rate for Early (n=26 neurons) and Late units (n= 18 neurons) around the time of induced CSM. Represents group mean and ± SEM across mice (N=3 mice). (**bottom**) Time of max firing rate difference relative to CSM onset time.

As in Fig. 2, we grouped IntA units based on their activity timing relative to reach – ‘Early Units’ peaked before, and ‘Late Units’ after, maximum outward velocity. To isolate stimulus- and CSM-related activity, we computed average firing rate changes for Stim_on_ relative to Stim_off_ (baseline) trials (***Fig. 5D***). Z-scored population averages revealed robust CSM-related modulation in both Early (n = 26) and Late (n = 18) Units (***Fig. 5C,*** middle/bottom). Surprisingly, Early Units were sometimes suppressed during the light pulse train (***Fig. 5B***, top, example data; **5C–D,** middle, blue box), but then showed a coordinated increase in activity aligned with the induced CSM (***Fig 5D***, middle, arrow, z=3.25, *p=0.0012*). In contrast, Late Units were consistently excited by optogenetic stimulation and exhibited a second activity peak during the late phase of the CSM (***Fig. 5B*** (bottom)***–D*,** bottom, arrow, z=2.56, *p=0.001*). Quantifying CSM-related firing across groups revealed robust rate increases – averaging 15 - 20 spikes/sec – during the CSM compared to unstimulated reaches (***Fig. 5E***, top). These increases occurred within milliseconds of CSM onset: Early Units peaked <5 ms before CSM initiation, while Late units peaked ∼25 ms afterward (***Fig. 5E***, bottom). Together, these findings show that IntA neurons are strongly recruited during induced CSMs and preserve their temporal dynamics – Early or Late – from the primary reach, now embedded within the structure of the induced CSMs.

### Behavioral and neural adaptation to repeated perturbation

We were ultimately interested in testing whether neural adaptation occurs in cerebellar output neurons following perturbations to reaches. Given that CSMs were instructive, leading to probabilistic adjustments in CSM timing, (***Fig. 4***), we designed an experiment to augment learning, introducing closed-loop perturbations of the IntA in a block format ^35^. We used a perturbation schedule in which mice made 15-25 unstimulated reaches (baseline block) followed by 25 reaches in which IntA^RN^ neurons were stimulated for 50 ms on every reach (stimulation block), concluding with a ‘washout block’ in which no stimuli were applied (***Fig. 6A***), and aligned data to the time of the induced CSM as the hypothesized instructive signal.

**Figure 6:**
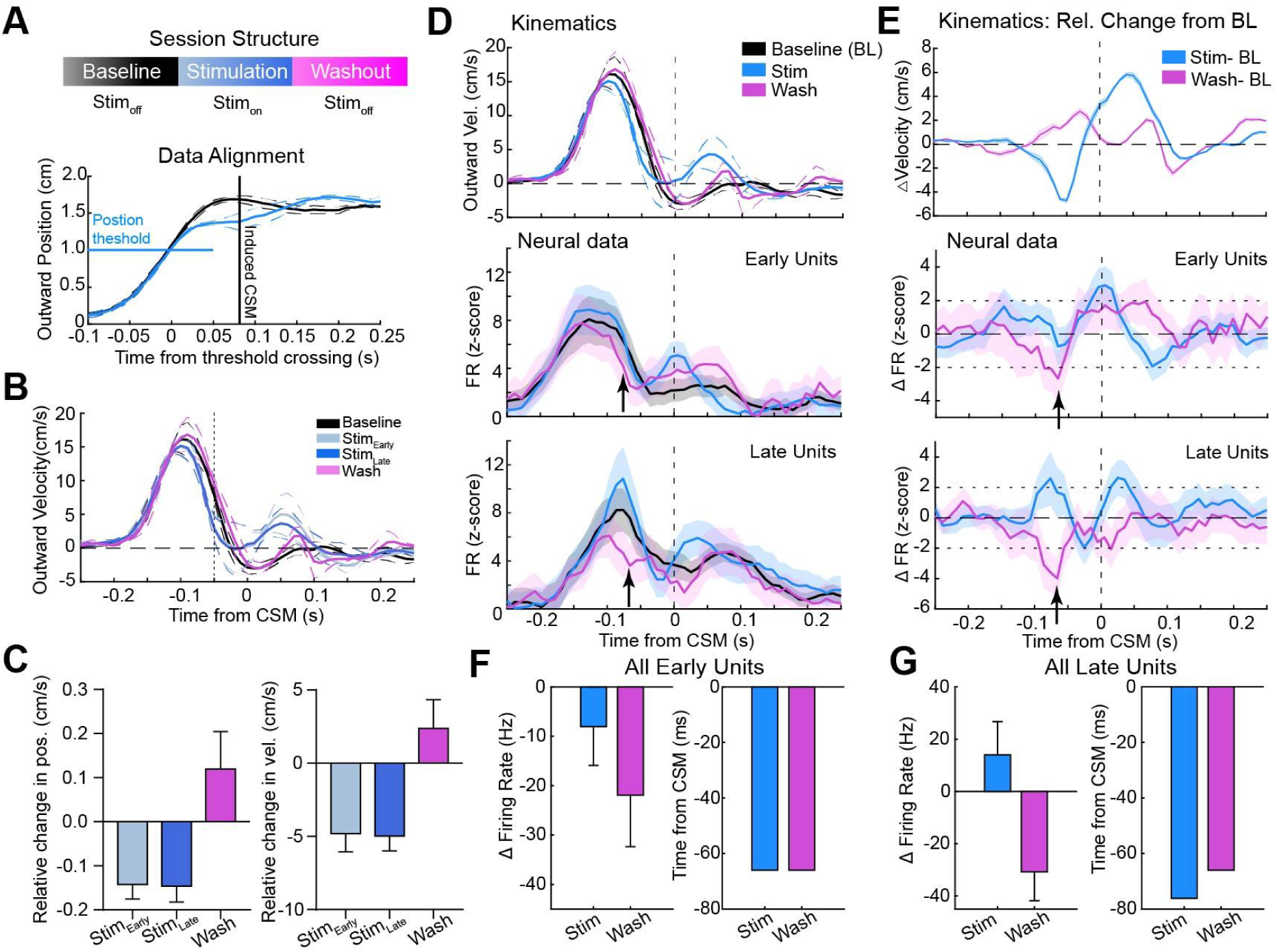
Behavioral and neural adaptation to repeated perturbation. (A) Session structure and data alignment. Behavioral sessions consisted of Baseline (black), Stimulation (blue), and Washout (magenta) epochs. Reach trajectories were aligned to the time of threshold crossing (0 s, vertical line) or to the induced CSM time (80 ms). (B) Velocity traces aligned to CSM onset (thick lines mean ± SE thin lines) for Baseline, Stimulation (Early and Late-lines overlap), and Washout conditions. The dashed line marks the analysis window used for computing changes in velocity and position, 50 ms before CSM onset. (C) Relative changes in position (left) and velocity (right) across Stimulation (Early, Late) and Washout conditions compared to Baseline. Error bars represent SEM. (D) Induced CSM-aligned velocity and firing rate. (**Top**) Mean outward velocity traces aligned to CSM onset. (**Middle** and **bottom**) Population firing rate (FR, z-scored) for Early Units (n = 26 neurons) and Late Units (n = 18 neurons), respectively. Shaded areas indicate SEM, N=3 mice. (E) Relative changes in velocity and firing rate. (**Top**) Velocity difference relative to Baseline for Stimulation and Washout. (**Middle** and **bottom**) Changes in firing rate (ΔFR, z-score) relative to Baseline for Early and Late Units. Middle plot: arrow, z = −2.67, *p=0.008*. Bottom plot: arrow, z= −3.98, p=<0.0001 (two-tailed z-test). Thin lines show mean change ± SEM. Small horizontal dashed lines indicate ±2 z-scores, corresponding to approximately the 95% confidence interval under a standard normal distribution (F–G) Summary of neural modulation. Group-level changes in firing rate (ΔFR, Hz, left) and CSM timing (right) during Stimulation and Washout for Early (F, n=26 neurons) and late (G, n=18 neurons) unit populations. Error bars represent SEM, N=3 mice.

As described above (***Fig. 5***), IntA^RN^ stimulation acutely reduced outward reach velocity (***Fig. 6 B-C***). Notably, across the block of stimulated trials, the magnitude of the perturbation effect remained stable (**Supplemental Fig. 5**), contrasting this manipulation of cerebellar outputs with the rapid adaptation to manipulation of cerebellar inputs from pontine mossy fibers described previously ^35^. Importantly, however, upon removal of stimulation during the washout block, we observed kinematic adaptation as subtle but consistent increases in outward position and velocity relative to baseline reaches, in the direction opposite of the stimulation effect, with the limb moving on average 2.4 cm/s faster than baseline reaches preceding CSM onset (***Fig. 6C, D-E*** *top*). Summarizing, these results show kinematic adaptation to optogenetic stimulation of IntA output, which was only visible during washout. Thus, although the rest of the brain may participate in generating real-time corrections that mitigate the behavioral consequences of the perturbation, the washout effect in the absence of a stimulation-block effect suggest that covert learning occurs upstream of this perturbation, presumably in PCs, expression of which was blocked by the optogenetic stimulation of cerebellar outputs.

This hypothesis predicts changes in cerebellar output during the washout block relative to baseline that would counter the kinematic perturbation. Consistent with this prediction, we observed a temporally restricted firing rate suppression in IntA population activity during the washout block, averaging 20-30 spikes/second below baseline (***Fig. 6D-E***, middle, bottom). This suppression was observed in both ‘Early’ and ‘Late’ IntA populations, and preceded the induced CSMs from the stimulation block by approximately 65 ms (***Fig. 6E***, middle, arrow z=-2.67, p=0.008; bottom, arrow z=-3.98, p<0.0001, two-tailed z-test). This timing coincided with the firing rate changes observed during the stimulation block and the stimulation-to-induced-CSM latency (***Fig. 6F-G***). We quantified these effects by measuring the largest magnitude rate changes relative to baseline, comparing the optogenetic stimulation block and the washout block, before the CSM epoch. Both Early and Late Units showed significant learned suppression during washout (***Fig. 6F-G***). This neural signature of adaptation in washout suggests that learning occurred upstream of the cerebellar nuclei, most likely within the cerebellar cortex. The direction of change (suppression) further suggests that parallel fiber-to-PC synapses underwent LTP, increasing firing rates during the perturbation block, at the approximate time of the perturbation. Given that a substantial population of IntA neurons exert an inward pull that decelerates outward limb velocity, our findings show that, through learning, IntA activity can be suppressed to reduce this decelerative drive when deceleration is too rapid because of the optogenetic effect, a behaviorally-relevant adaptation expressed through cerebellar output^32,35,47–51^.

## Discussion

We found that corrective sub-movements are not only common during naturalistic reach-to-grasp movements in mice, but also instruct learning. Mice generate CSMs that counter both spontaneous and induced deviations from average reach kinematics. IntA neuronal activity was correlated with both spontaneous and induced CSMs, suggesting involvement in feedback corrections. However, the primary reach-related modulation of IntA activity was not related to CSMs, suggesting a predominant role in feedforward control of reach kinematics. Induced CSMs were instructive to the learned timing of subsequent spontaneous CSMs and drove neural adaptation that opposed optogenetically induced perturbations. Adaptation was nevertheless covert – learning expression was blocked by the perturbation but unmasked in the washout block. This suggests a site of learning upstream of IntA, likely involving PCs^52–54^. In aggregate, our results show neural expression of adaptation in the cerebellar nuclei and link this form of anticipatory ‘feedforward’ control with the expression of feedback-driven error corrections.

Work in humans and non-human primates (NHPs) has identified CSMs as a hallmark of optimal feedback control, enabling real-time corrections to motor output ^7,25,27,28^. In our analysis, CSMs were consistently preceded by trajectory deviations, supporting the idea that they function as corrective adjustments rather than as random motor noise ^4,55^. Interestingly, in mice naive to perturbations, spontaneous CSMs occurred throughout the reach trajectory, but their probability was not uniform; instead, they showed a multimodal distribution, suggesting intermittent control. Similar midflight corrections with intermittent enhanced probability have been observed in humans and NHPs ^42,43,56–59^. Given the proposed instructive role of CSMs, we speculate that they may be associated with complex spikes, either increases or decreases driving plasticity in PCs. Such distributed, non-uniform feedback could help maintain continuous, learned reach trajectories by segmentally reinforcing neural patterning through plasticity at parallel fiber–PC synapses. Moreover, such a process could be supported by reach-phase specialized neurons, such as early and late cells, that control distinct temporal phases of reach.

Rapid error correction depends on intact motor cortex and spinal circuits, both for goal-targeting errors and responses to acute perturbations^8,60^. The fastest responses are mediated by short-latency spinal reflexes, reflecting central predictions that regulate stretch reflex gain via gamma motor neurons. In contrast, longer-latency corrections involve transcortical pathways ^61–63^. For example, cortical inactivation in mice impairs corrective movements after target undershoots and disrupts responses to induced perturbations ^4,5,64^. Similarly, online corrections in tongue movements require motor cortical activity, and re-aiming after spout misses depends on the superior colliculus ^4,65–67^. Given these findings, how might cerebellar signaling contribute? Previous work in monkeys demonstrated that cooling cerebellar output during reach perturbations did not eliminate rapid corrections but delayed their stopping, resulting in overcorrections ^68–70^. Based on this, it was proposed that under normal conditions, motor cortex generates a braking response to terminate reflex-driven corrections, with cerebellar input providing the predictive signal that times this braking ^18^. Our data offer a refinement of this interpretation. Specifically, we suggest that the cerebellum, via IntA activity during the late phase of the CSM, may itself generate the predictive braking signal – not merely supply predictive information to motor cortex. Disrupting cerebellar output would therefore eliminate this predictive brake, leading to poorly timed and/or scaled corrections. These cerebellar signals may act through ascending pathways to cortex ^64^ or bypass cortical circuits entirely via descending projections through the brainstem or directly to the spinal cord ^33,71–73^. Future studies targeting diverse push-pull modules within the cerebellar nuclei could indicate even more intricate agonist-antagonist coordination of CSMs by cerebellar output.

In many human and NHP studies of reach adaptation, CSMs are experimentally evoked by applying a forcefield or jumping the target ^57,74–77^. We found CSMs in response to brief closed-loop optogenetic stimulation of IntA^RN^ neurons. Notably, when we used a complementary manipulation, stimulating both the excitatory and inhibitory neurons of IntA^All^, CSMs were inaccurate and lacked temporal stereotypy. These CSMs also did not drive changes in spontaneous CSM timing on subsequent reaches. These results raise the question as to why CSMs were delayed and learning was disrupted. Intriguingly, because these manipulations were non cell-type specific, they likely included manipulation of inhibitory neurons that project to the inferior olive. Thus, we may have disrupted the ability of the system to remain sensitive to error and/or drive learning as a result of the response to error. These findings support a model in which the cerebellum contributes to real-time movement execution via nuclear output, acting primarily as a feedforward controller but with roles in speeding and learning from feedback control. Moreover, these results provide a plausible mechanism by which feedforward control is influenced by feedback corrections and vice versa, a recursive relationship identified in humans and cerebellar patients ^15–17,78,79^.

In a previous study, we found that repeated stimulation of cerebellar mossy fiber inputs perturbed reaches acutely, but the effect of stimulation rapidly diminished ^35^, similar to recently described gain adaptation in mouse forelimb movements ^80^. When the perturbation was omitted, opposite-direction after-effects revealed adaptation. Here, we used the same trial structure but targeted cerebellar outputs, rather than inputs. Unlike with input perturbations, reaches failed to compensate for the stimulation during the adaptation block. However, upon removal of stimulation, subtle behavioral aftereffects and opposite-direction neural adaptation in IntA firing was observed. Specifically, a strong suppression of IntA firing approximately 65 ms before the induced CSMs emerged during the washout block, in the opposite direction as the optogenetic activation. This time delay is approximately the error:CSM latency, suggesting the potential for delayed credit-assignment to synaptic inputs to PCs ^81^. An attractive explanation is that the suppression of IntA activity is caused by increased PC activity from induction of long-term potentiation (LTP) at parallel fiber-PC synapses, predicting that these perturbations lead to parallel fiber activity but reduced climbing fiber firing in this module. However, alternative or complementary mechanisms may also contribute. These include homeostatic plasticity within IntA neurons in response to sustained artificial activation, short-term or long-term depression of mossy fiber collateral input to the IntA, or disrupted error signaling from the inferior olive due to altered movement feedback.

Together, these findings suggest that the cerebellum can still engage plasticity mechanisms under conditions of output disruption; however, the brain as a whole does not compensate for disruption of cerebellar output on these time scales. These rules may explain why extra-cerebellar motor learning systems do not fully mitigate erroneous cerebellar output in disease ^82–84^. In such cases, feedback corrections mediated by other brain regions may exhibit lag ^4,6,85,86^ by relying on sensory feedback in the absence of predictive cerebellar signals, contributing to phenotypes observed following cerebellar damage.

## SUPPLEMENTAL FIGURES

**Supplemental Figure 1.**
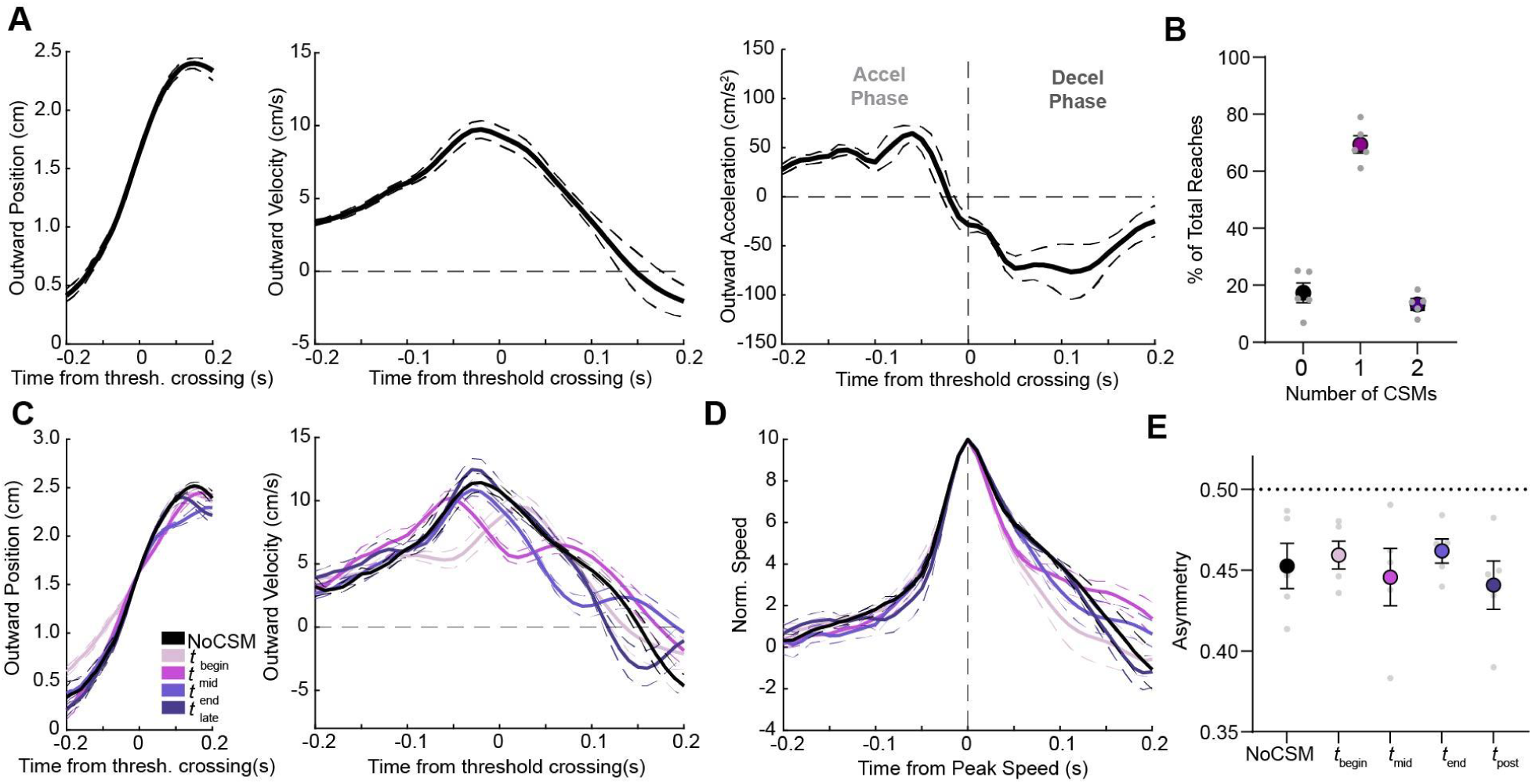
Additional kinematic characterization of reaches with and without CSMs. (A) Group-averaged outward position (left), velocity (middle), and acceleration (right) trajectories aligned to threshold crossing (time = 0) for Sham mice (N = 5 mice, n=2074 reaches). Gray vertical line separates acceleration and deceleration phases. Thin dashed lines: ±SE ; bold lines: mean across mice. (B) Distribution of reaches containing 0, 1, or 2 CSMs, shown as a percentage of total reaches across mice. Gray dots represent individual mice; black and purple bars denote group means ± SEM. (C) Outward position (left), velocity (middle), and acceleration (right) traces aligned to threshold crossing and separated by CSM timing (NoCSM in black; early to post-CSM in increasing shades of light purple/dark purple). Thin dashed lines: ±SE ; bold lines: mean across mice. (D) Average normalized speed profiles aligned to peak speed for each CSM timing group. Thin dashed lines: ±SE ; bold lines: mean across mice. (E) Reach asymmetry index : asymmetry calculated by dividing the area under the curve (AUC) of the first half of the speed profile by the total AUC. Dotted line at 0.5 denotes symmetric reaches. All reach categories have varying levels of asymmetry. Points show individual mice; bars indicate mean ± SEM. (RM One-Way ANOVA, *p* = 0.65, *N* = 5 mice).

**Supplemental Figure 2:**
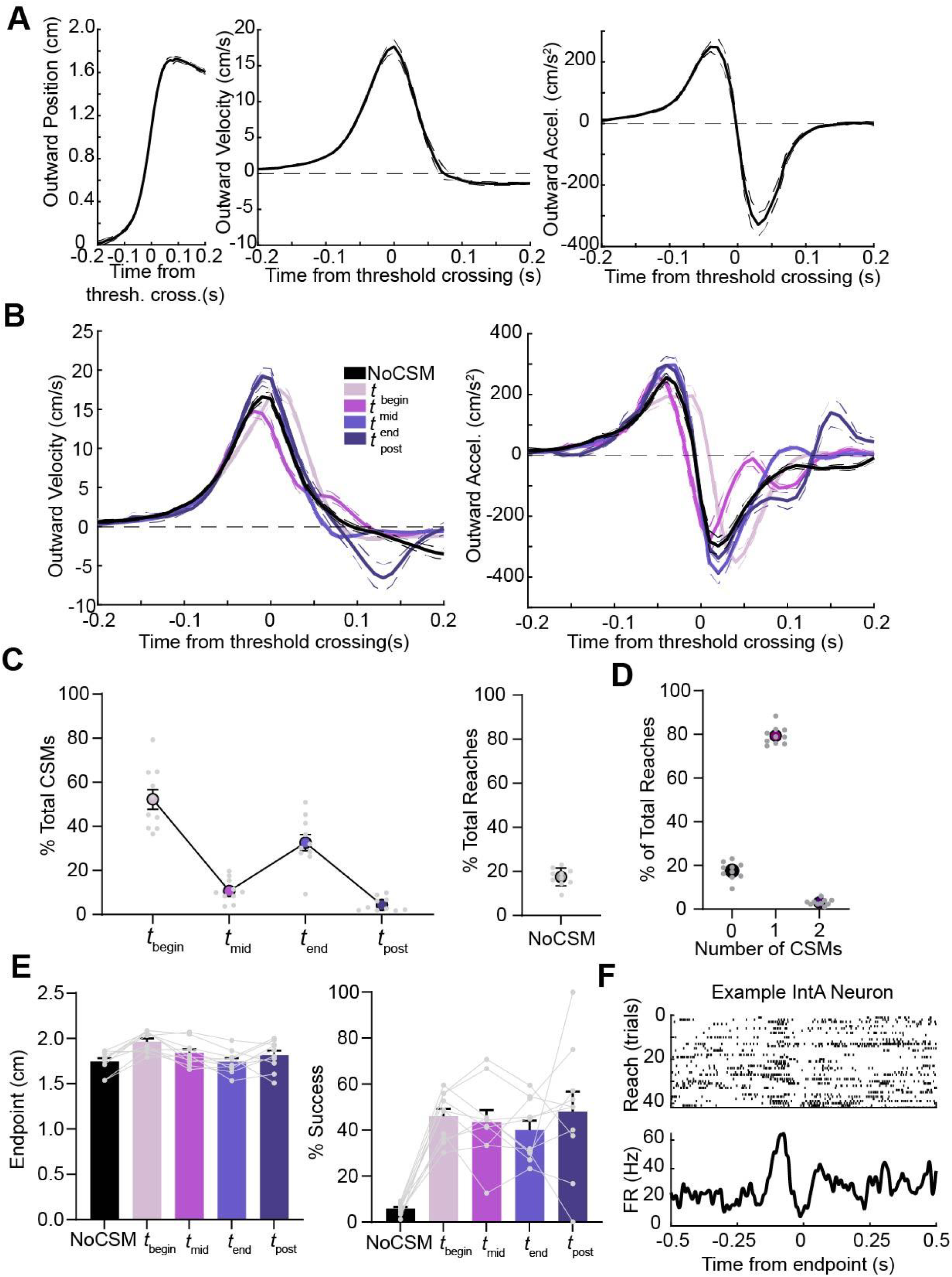
Head-fixed reaching CSM characterization. (A) Group-averaged outward position (left), velocity (middle), and acceleration (right) trajectories aligned to threshold crossing (time = 0) for Head-fixed Sham mice (N = 10 mice). Thin dashed lines: ±SE ; bold lines: mean across mice. (B) Outward velocity (left) and acceleration (right) traces aligned to threshold crossing and separated by CSM timing (NoCSM in black; beginning to post-CSM in increasing shades of light purple/dark purple). Thin dashed lines: ±SE ; bold lines: mean across mice. (C) Percentage of total reaches containing CSMs within each epoch. Reaches without CSMs are shown for reference. Individual mice, gray circles; averages across mice, large dots (±SEM). (D) Distribution of reaches containing 0, 1, or 2 CSMs, shown as a percentage of total reaches across mice. Gray dots represent individual mice; black and purple bars denote group means ± SEM. (E) Left: Average endpoint position for reaches with No CSM and reaches with begin, mid, end, or post CSMs (RM One-Way ANOVA, *p* = 0.002, Post-Hoc Comparisons (Tukey) NoCSM and t_begin_ (*p* = 0.002) and between t_begin_ and t_end_ (*p* < 0.0001) *n* = 10 mice). Right: Proportion of successful reaches across conditions (RM One-Way ANOVA, *p* = 0.11, Post-Hoc Comparisons (Tukey); No CSM: t_begin_ (*p* < 0.0001), t_mid_(*p* = 0.0004), t_end_ (*p* = 0.0001), and t_post_ (*p* = 0.008) *n* = 10 mice). Light gray circles: individual mice ; bars: group mean ± SEM. (F) Raster plot (top) and peristimulus time histogram (PSTH; bottom) from an example IntA neuron aligned to reach endpoint.

**Supplement Figure 3:**
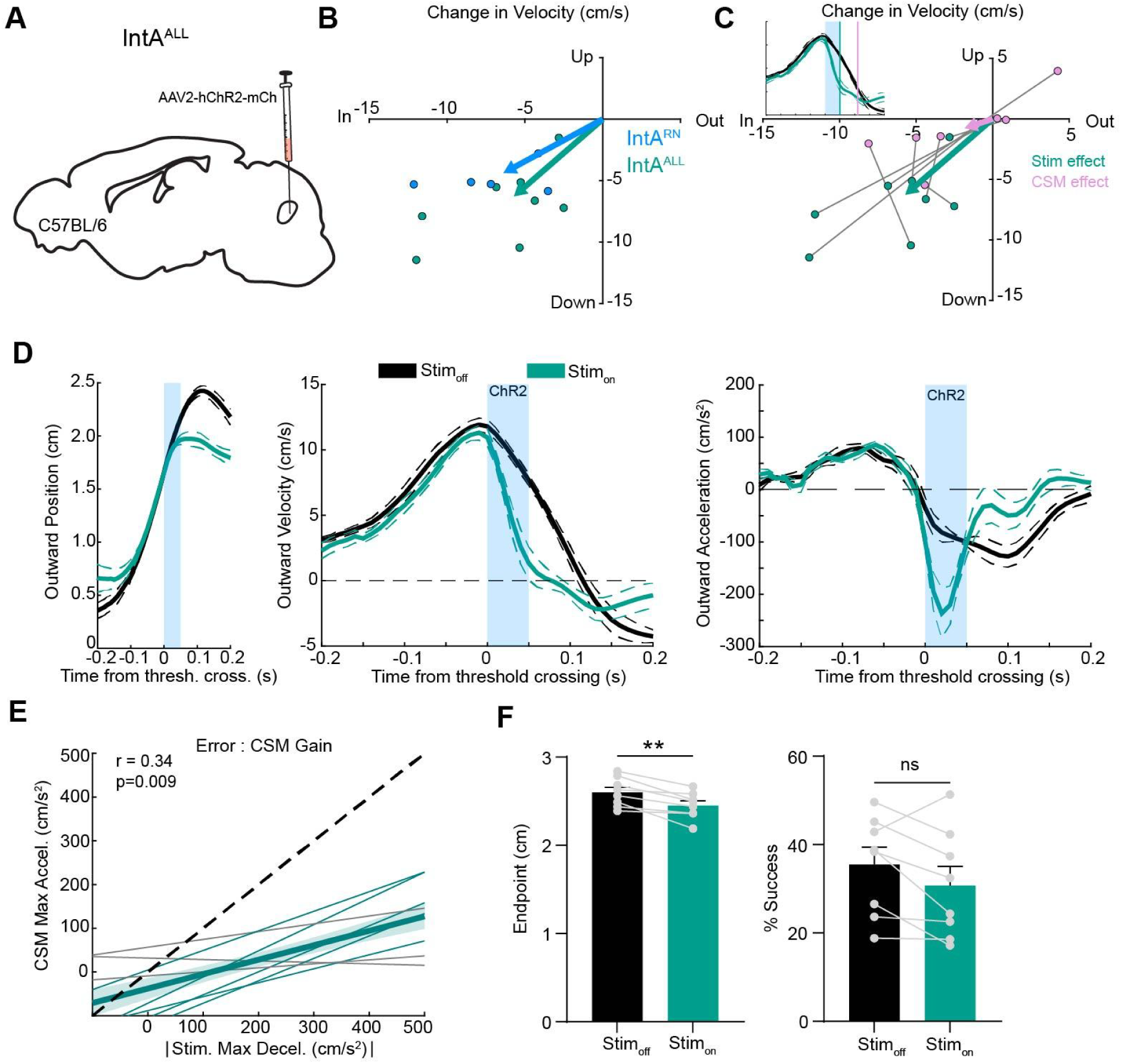
Optogenetic perturbation of all IntA output during reaching disrupts ongoing kinematics. (A) Schematic of viral targeting strategy. AAV2-hChR2-mCh was injected unilaterally into the anterior interposed nucleus (IntA) of C57BL/6 mice. (6 of 8 C57/Bl6 mice comprised a dataset used in Becker and Person, 2019, reanalyzed here). (B) Average change in outward and upward velocity for each animal at the (Stim_on_ - Stim_off_) for IntA^RN^ and IntA^All^ experiments. The average effect size is indicated by arrow length (IntA^RN^ N=5 mice, *p=0.003*, (outward, −7.09 ± 1.56 cm/s, upward, −4.87 ± 0.54 cm/s), IntA^All^ N=8 mice, *p=0.0004*, one sample t test (outward, −6.34 ± 1.29 cm/s, upward −6.98 ± 1.11 cm/s)). Comparison between groups (Outward: *p=0.72*, Upward *p=0.18*, two sample t-test). (C) Average change in outward and upward velocity for each animal at the (Stim_on_ - Stim_off_) for 50ms after (Stim effect) and 110 ms after threshold crossing (CSM effect). The average effect size is indicated by arrow length (Stim effect, *p=0.0004*, one sample t test (outward, −6.34 ± 1.29 cm/s, upward −6.98 ± 1.11 cm/s), CSM effect, *p=0.008*, one sample t test (outward, −2.04± 1.39 cm/s, upward −0.86 ± 0.93 cm/s)). Comparison between groups (Outward: *p=0.04*, Upward *p=0.0008*, two sample t-test). N=8 mice. (D) Kinematics aligned to threshold crossing (Stim, blue shaded region): outward position (left), velocity (middle), and acceleration (right); (mean ± SE; N = 8 mice). Traces compare trials with and without stimulation (Stim_off_, black; Stim_on_, teal). (E) Across mice, greater optogenetically induced deceleration (x-axis: max outward deceleration during stimulation epoch) was weakly associated with compensatory CSM responses (y-axis: max outward acceleration during CSM) (r = 0.34, p = 0.009, Fisher z t-test, N=8 mice, only 5 out of 8 animals had significant correlations). Individual lines represent the mean of each animal, and the thick line represents the group mean ± SEM across mice. The dashed line indicates unity. (F) Endpoint position (left) (p=0.004). Success rate dropped in 5 of 8 mice although as a population this was non-significant (right) (p = 0.1, paired t-test). Gray lines show matched data per animal, N=8 mice.

**Supplemental Figure 4:**
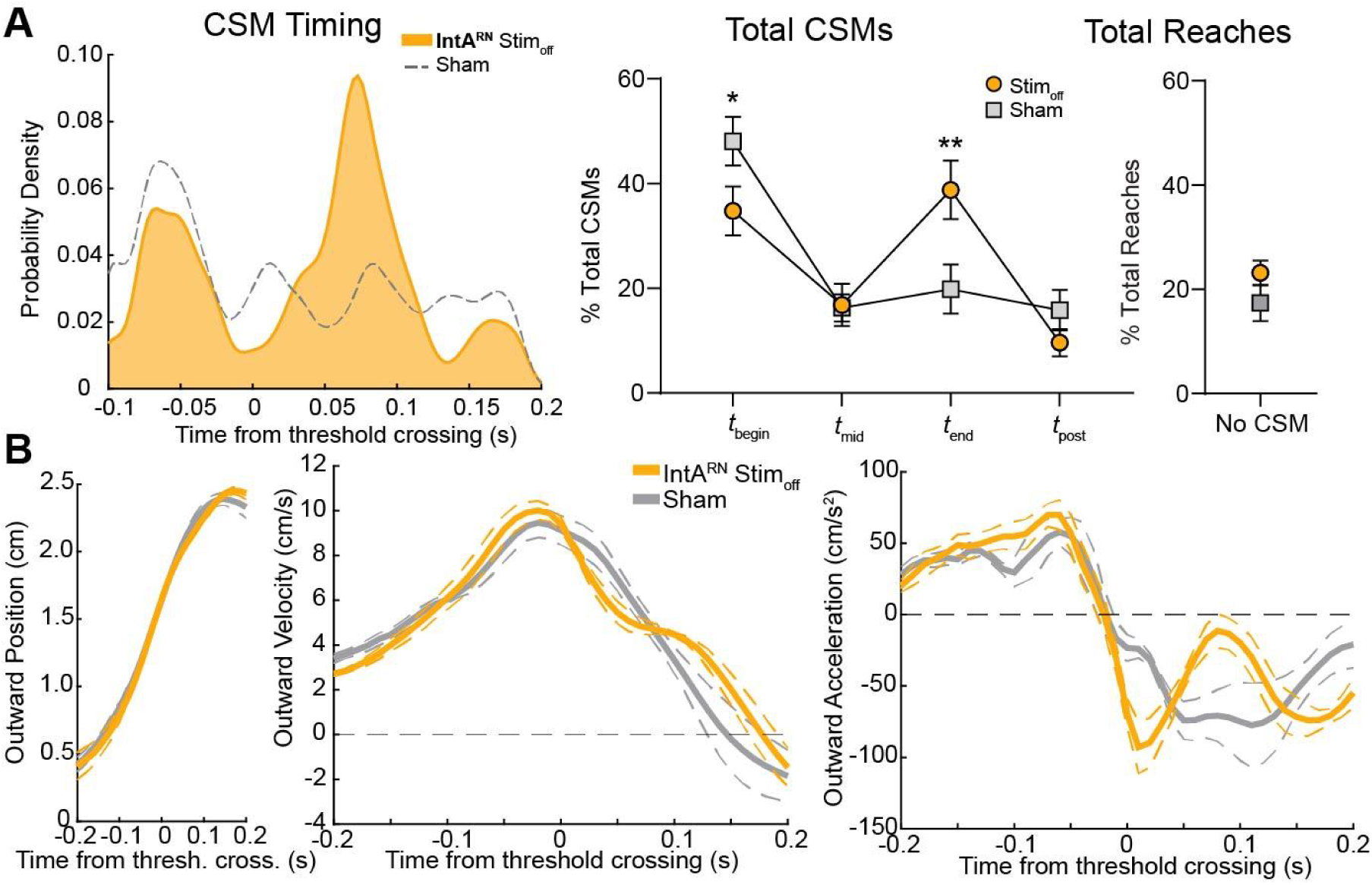
Direct comparison of IntA^RN^ Stim_off_ reaches to Sham condition. (A) CSM timing (p<0.001, D=0.17, Kolmogorov–Smirnov test),Total CSMs(p=0.004, two-way ANOVA, multiple comparisons: t_begin_ p=0.03, t_end_ p=0.003), Total Reaches containing CSMs (p=0.17, paired t-test). IntA^RN^ N=5 mice; Sham N= 5 mice. (B) Kinematics aligned to threshold crossing: position (left), outward velocity (middle), and acceleration (right); (mean ± SE). Traces compare trials from animals that received optogenetic stimulation (IntA^RN^ N=5 mice) to those that did not (Sham N= 5 mice).

**Supplemental Figure 5:**
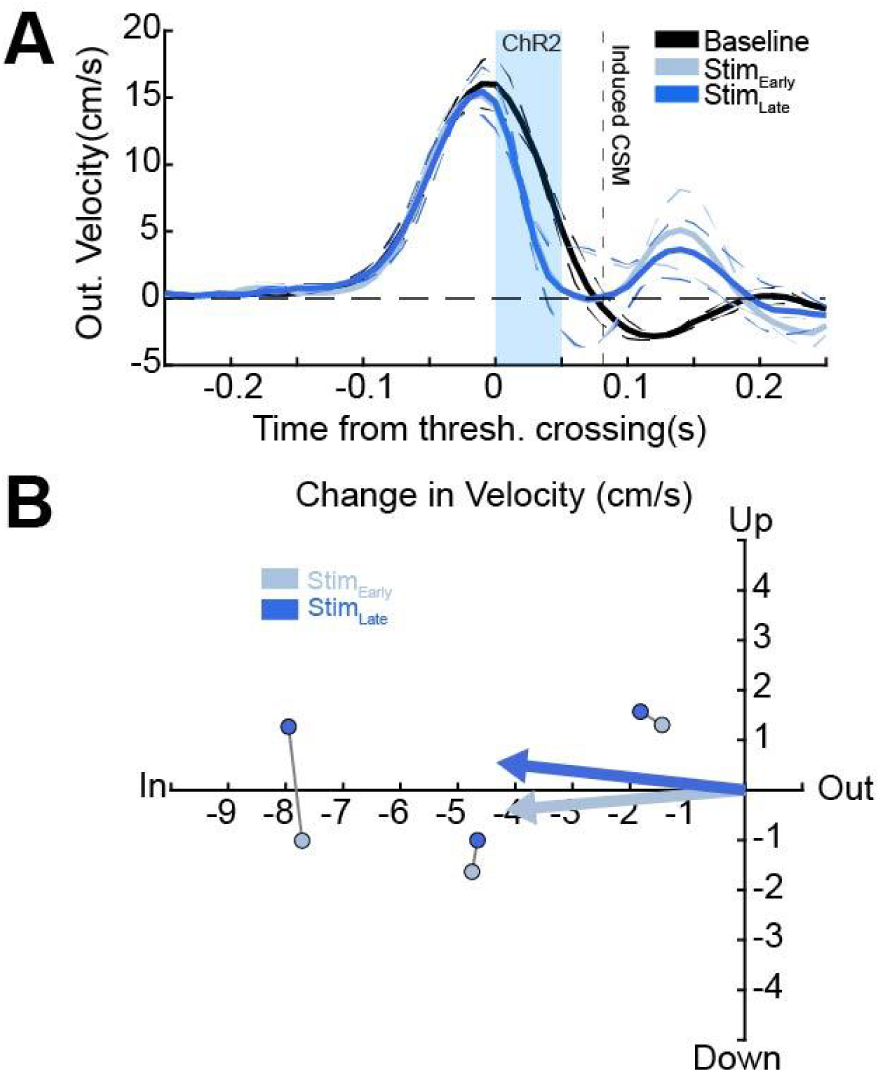
No overt behavioral adaptation during IntA^RN^ block stimulation. (A) Velocity traces aligned to time from threshold crossing (thick lines mean ± SE thin lines) for Baseline, Stimulation (Early and Late). The dashed line marks the timing of the induced CSM. (B) Average change in outward and upward velocity for each animal at the end of the stimulation epoch (Stimon - Stim_off_) for the first 5 reaches during the stimulation block and the last 5 reaches of the stimulation block. The average effect size is indicated by arrow length (Stim_Early_, *p=0.1*, one sample t-test (outward, −4.6 ± 1.8cm/s, upward, −0.5 ± 0.9 cm/s); Stim_Late_, *p=0.09*, one sample t test (outward, −4.8 ± 1.8 cm/s, upward 0.6 ± 0.8 cm/s)). Comparison between groups (Outward: *p=0.95*, Upward *p=0.43*, two sample t-test). N=3 mice.

## STAR METHODS

### Animals (Experimental model and subject details)

Animals were housed in an environmentally controlled room, kept on a 12h light– dark cycle with ad libitum access to food and water, except during behavioral training and testing as described below. Behavioral and electrophysiology data were collected from 37 adult mice (P70-P365) of either sex (17 males, 20 females) from the following genotypes: (1) wildtype C57BL/6 (N=8); (2) Vglut2-cre (Vglut2-ires-Cre, Strain #016963, The Jackson Laboratory, N=29). After surgery, mice were individually housed with a running wheel. IntA^All^ analyses were performed on datasets used in Becker and Person, 2019. All procedures followed National Institutes of Health Guidelines and were approved by the Institutional Animal Care and Use Committee at the University of Colorado Anschutz Medical Campus.

### Surgical procedures and histology

All surgical procedures were conducted under ketamine-xylazine anesthesia. After induction of anesthesia, the surgical site was sterilized and bupivacaine was injected subcutaneously (2.5 mg/ml). Pressure viral injections were performed using a 10ul Hamilton Syringe, UMP3 microinjection syringe pump, and pulled glass pipette. The stereotaxic location of anterior interposed nucleus (IntA) was targeted as −1.9 mm posterior, 1.6 mm lateral, and −2.1 mm ventral from lambda. The stereotaxic location of the red nucleus (RN) was targeted as −3.5 mm posterior, −0.5 mm lateral and −3.6 ventral from bregma. For IntA^All^ experiments, we injected AAV2-hSyn-hChR2 (H134R)-mCherry in the IntA; for IntA^RN^ experiments we injected AAVretro-EF1a-Flpo in the RN and AAV8-hSyn-ConFon hChR2(H134R)-EYFP in the IntA; for head-fixed reaching and recording experiments we used AAV2-EF1α.DIO.hChR2 (H134R)-mCherry. Approximately 150 nL of virus was injected unilaterally into the right IntA and 150 nL into the contralateral left RN. For experiments in freely behaving mice, optical fibers (105 um core diameter, ThorLabs) were attached to a ceramic ferrule (1.25mm, Thor Labs), polished to >80% efficiency. They were then inserted 0.1 mm above the (−2.0 mm ventral) IntA and affixed to the skull using luting (3M) and dental acrylic (Teet’s cold cure). For Sham experiments, the same experimental protocol as IntA^RN^ experiments was followed; however, mice did not have viral expression and had no stimulation effect. For experiments in head-fixed mice, an optical fiber targeted the RN and was affixed to the skull along with a custom made aluminum head plate, centered on bregma. To allow for adequate viral expression time, experiments began 4-6 weeks following surgery. After experiments were completed, mice were euthanized according to standard procedures via pentobarbital overdose, transcardially perfused with 4% paraformaldehyde and processed for histological analysis. Injection sites and fiber implant tracts were verified using an upright epifluorescent microscope.

### Reach behavior

Following recovery from surgery, mice were trained on a skilled reach task in which they were food restricted to 80-90% of their baseline weight for reach training. Mice were monitored daily to ensure proper weight loss. They were then habituated to the behavioral arena or the head-fixed apparatus by presenting food pellets on a pedestal (20 mg, BioServ #F0163) that could be reached by their tongue. The head-fixed reaching apparatus was adapted from Mosberger et al. 2024, in which mice are positioned in a cup shaped holder that allows for a wider range of forelimb movements ^87^. Once mice started eating from the pedestal, the pedestal was slowly moved to encourage reaching with the right forelimb. Training sessions lasted until mice were actively engaged and reaching for 20-30 minutes. Mice were trained for a minimum of 1 week before being considered fully trained and once they could successfully retrieve 40% of pellets for 3 days in a row.

### Kinematic Tracking and closed loop optogenetic stimulation

Closed loop kinematic tracking was conducted as previously detailed ^32,35^. We used a 50 ms pulse train at 100 Hz, 2 ms pulse width to activate ChR2 (IntA^RN^, IntA^All^) with a 470 nm diode laser (Opto Engine LLC), for both randomized and block stimulation paradigms. For randomized sessions, stimulation was delivered on a pseudorandom 25% of trials. In block stimulation experiments, sessions followed a structured format: a **Baseline** block of 15–25 unstimulated reaches followed by a **Stimulation** block of 20–25 consecutive reaches with stimulation on every reach, and finally a **Washout** block with no stimulation that lasted for as long as the mouse continued to engage in the task. The strategy used in block stimulation experiments targeted optical activation of IntA terminals in the RN, allowing for specific targeting of the IntA-RN pathway and mitigated optoelectric artifacts.

Laser output was monitored and calibrated before each session, with power ranging from 0.5-2 mW. Laser output varied individually between mice depending on the stimulation effect. For freely behaving experiments, stimulation was delivered at the positional threshold of 1.6 cm from the origin point inside the behavior box, while for the head-fixed experiments it occurred at the positional threshold of 1.0 cm from the origin point at the perch. Both threshold points are near the average position of peak outward velocity for their respective conditions ^32,35^.

### Kinematic analysis

All initial processing of raw kinematic data was performed using custom written MATLAB scripts as previously described ^32,35^. Corrective submovements (CSMs) were identified as deviations in acceleration during outward reaches. By aligning reaches to a positional threshold, we could examine differences in peak velocity or timecourse, relative to a consistent landmark; additionally, the alignment threshold was the positional trigger for closed-loop stimulation, thus we could also reference non-stimulated and stimulated trials similarly using this alignment. CSMs were operationally defined as deviations in the acceleration or jerk profiles. During the deceleration phase of the reach, CSMs were identified within a 200 ms window centered on a mid-reach positional landmark, referred to as the “threshold”. A trial was classified as containing a CSM if there was a positive reacceleration, identified by a zero-crossing in the acceleration trace and if the reacceleration was sustained for more than 25 ms. For reaches in which the CSM occurred before threshold crossing, we instead identified the position of maximum jerk, indicating a rapid increase in acceleration. All CSMs were visually inspected across mice and sessions to ensure robust detection and to exclude false positives due to noise or tracking artifacts. Finally, behavioral kinematics (position, velocity, and acceleration) were averaged and visualized to reveal characteristic movement patterns associated with early corrections.

We stratified CSM-containing reaches by CSM time within the reach, illustrating the grand mean across of these groups of mice. Four distinct modes were identified in the grand mean distribution of CSM times across mice. Temporal boundaries were defined between adjacent local minima surrounding each peak (Freely Behaving: Distribution peaks were clustered in early (t_begin_ : −0.1 - 0 s), mid (t_mid_ : 0 - 0.059 s), late (t_end_: 0.06 - 0.132 s)), and post (t_post_: t <0.133 s) epochs). Given the differences between the distributions between CSMs in the freely behaving versus head-fixed condition, distribution peaks were shifted slightly for head-fixed reaching (Head-fixed: Distribution peaks were clustered in early (t_begin_ : −0.1 - 0 s), mid (t_mid_ : 0 - 0.06 s), late (t_end_: 0.06 - 0.12 s)), and post (t_post_: t <0.12 s) epochs). For kinematic alignments to CSM epochs, we used the average CSM time within each epoch. For the sliding z-score analyses of kinematics, all reaches from each animal were aligned to the relevant epoch and averaged to generate a “template reach.” These templates were then compared to subsets of reaches with CSMs within particular epochs. For each animal, we computed point-wise z-scores by normalizing the CSM-epoch-sorted reaches to the mean and standard deviation of the corresponding template reaches. We then averaged normalized traces across mice. For the block stimulation experiments, early and late stimulation reaches were defined as the first five and last five reaches within each block, respectively. For the washout condition, the first five reaches were analyzed to capture immediate learning effects ^35,80^.

### Neuropixels recordings

In preparation for recordings, craniotomies were made over the IntA, ipsilateral to the reaching arm, in fully trained mice. The brain was covered in sterile saline and sealed with Kwik-Sil silicone to preserve the recording site for multiple acute recordings. Recordings were made using Neuropixels 1.0 probes. Probes were lowered into the brain using a motorized micromanipulator (Sensapex uMp micromanipulator). Once the electrode shank passed the putative purkinje cell layer and entered the nuclear layer, we waited 15 minutes before starting a baseline recording of 10 minutes, then began the behavioral recordings. Electrophysiology data was acquired using an OpenEphys system (https://open-ephys.org/gui). Data was sorted offline in Kilosort2 and manually curated in phy (https://github.com/cortex-lab/phy). 3-10 recordings were made from each animal. After each recording, the craniotomy was cleaned and filled with silicone gel. For the final recording, we coated the Neuropixels probe with DiI to facilitate post-hoc histological identification of the recording site.

### Neural data analysis

Following sorting, units were analyzed offline using custom written Matlab code ^35^. IntA neurons were identified by depth on the Neuropixles probe as well as whether or not they were modulated by reach. To assess whether individual neurons were modulated during reach, we compared firing rates during reach related and non-reach related timepoints. For each neuron, we computed the mean firing rate during reach (FR_reach) and non-reach (FR_nonreach) periods and quantified modulation using a modulation index: (FRreach-Frnonreach)/(FRreach+FRnonreach).

A neuron was considered reach-modulated if its absolute modulation index exceeded 0.1. Additionally, we performed a Wilcoxon rank-sum test cell by cell to assess statistical differences in firing rates between reach and non-reach periods. This approach allowed us to systematically identify neurons whose activity was significantly modulated during reaching behavior.

We restricted further analyses to units that were active during the accelerative or decelerative phases of the reach. Units were classified based on whether their trial averaged peak firing rate occurred within 100 ms before (Early Units) or after (Late Units) peak velocity. For each unit, the raw PSTH was then smoothed using a Gaussian kernel with a standard deviation of 5 ms. Both raw and smoothed PSTHs were retained for subsequent analyses. For population level analyses, we computed the z-score of the firing rate of each unit relative to non-reach baseline, facilitating comparison across units with different baseline firing rates. The resulting z-scored activity was then averaged across the population to assess overall patterns of neural modulation. To quantify the magnitude of firing rate changes during optogenetically perturbed reaches, we analyzed smoothed PSTHs aligned to threshold crossing. For each group we identified the peak firing rate change following stimulation offset and calculated the corresponding SEM across units. This provided a summary measure of population level response magnitude of the corresponding induced CSM. The same approach was used to quantify learned changes in firing rate following the block stimulation protocol. In this case, both the behavioral and neural data were aligned to the time of the average induced CSM.

### Quantification and Statistical Analysis

Statistical analyses were performed using standard tests in MATLAB and GraphPad Prism. All data are displayed as mean ± SEM, unless otherwise noted in the text. All data were tested for normality using Kolmogrov-Smirnov test to choose between parametric and non-parametric statistical tests. Nonparametric tests were used when the dataset violated normality assumptions.All statistical tests and results are stated in the figure legends and text, along with population sizes for mice (N), reaches (n) and cells (n). Repeated Measure One-Way ANOVA with Geisser-Greenhouse correction and Tukey test used to correct for multiple comparisons as well as two-way ANOVA with Tukey test used to correct for multiple comparisons. The data reported in this paper reflect statistical summaries from each animal across multiple sessions. No statistical tests were used to predetermine required sample size. The experimenters were not blind to conditions during experiments and outcome assessments.

### Data and software availability

Available upon request to the corresponding author.

## Acknowledgements

We thank past and present members of the Person Lab for their feedback on this manuscript. Optogenetics support was provided by the University of Colorado Optogenetics and Neural Engineering Core. This work was supported by a National Research Service Award Individual Predoctoral Fellowship (F31) from NINDS (NS130867) to C.I.D. and NS131839 and NS114430 to A.L.P.

## Author Contributions

C.I.D. and A.L.P. designed and conceived the experiments. C.I.D. performed experiments and analyzed data, C.I.D. and A.L.P. interpreted results of the experiments. C.I.D. and A.L.P prepared figures and drafted and edited the manuscript. M.I.B. performed experiments and edited the manuscript.

## KEY RESOURCES TABLE

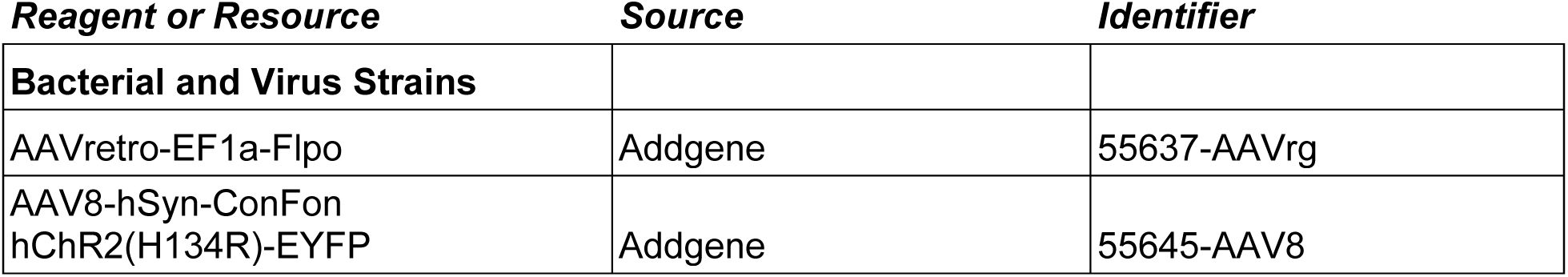
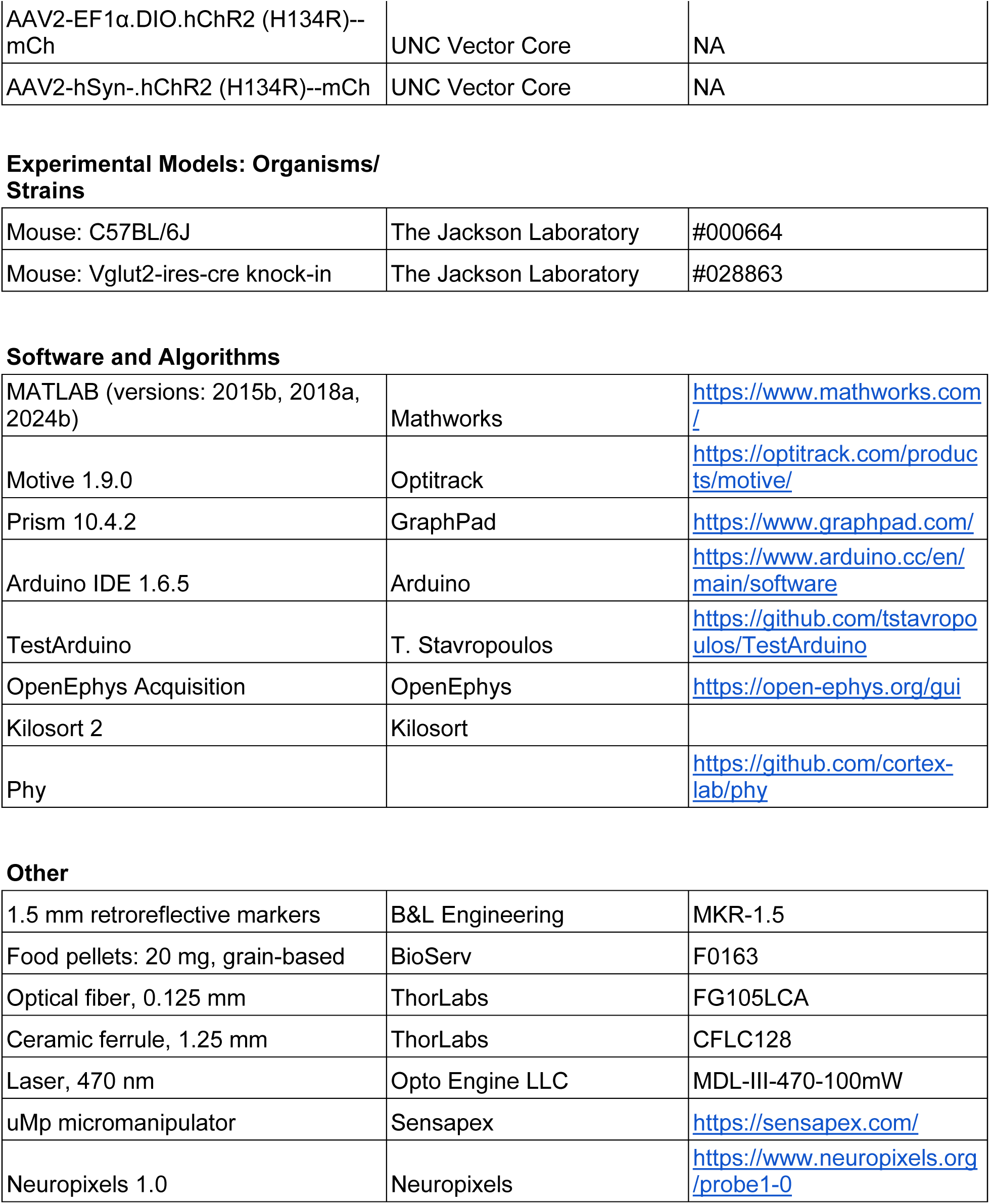

| ***Reagent or Resource*** | ***Source*** | ***Identifier*** |
| --- | --- | --- |
| **Bacterial and Virus Strains** |  |  |
| AAVretro-EF1a-Flpo | Addgene | 55637-AAVrg |
| AAV8-hSyn-ConFon<br>hChr2(H134R)-EYFP | Addgene | 55645-AAV8 |
| AAV2-EF1α.DIO.hChR2 (H134R)--mCh | UNC Vector Core | NA |
| AAV2-hSyn-.hChR2 (H134R)--mCh | UNC Vector Core | NA |
| Experimental Models: Organisms/Strains |  |  |
| Mouse: C57BL/6J | The Jackson Laboratory | #000664 |
| Mouse: Vglut2-ires-cre knock-in | The Jackson Laboratory | #028863 |
| Software and Algorithms |  |  |
| MATLAB (versions: 2015b, 2018a, 2024b) | Mathworks | https://www.mathworks.com/ |
| Motive 1.9.0 | Optitrack | https://optitrack.com/products/motive/ |
| Prism 10.4.2 | GraphPad | https://www.graphpad.com/ |
| Arduino IDE 1.6.5 | Arduino | https://www.arduino.cc/en/main/software |
| TestArduino | T. Stavropoulos | https://github.com/tstavropoulos/TestArduino |
| OpenEphys Acquisition | OpenEphys | https://open-ephys.org/gui |
| Kilosort 2 | Kilosort |  |
| Phy |  | https://github.com/cortex-lab/phy |
| Other |  |  |
| 1.5 mm retroreflective markers | B&L Engineering | MKR-1.5 |
| Food pellets: 20 mg, grain-based | BioServ | F0163 |
| Optical fiber, 0.125 mm | ThorLabs | FG105LCA |
| Ceramic ferrule, 1.25 mm | ThorLabs | CFLC128 |
| Laser, 470 nm | Opto Engine LLC | MDL-III-470-100mW |
| uMp micromanipulator | Sensapex | https://sensapex.com/ |
| Neuropixels 1.0 | Neuropixels | https://www.neuropixels.org/probe1-0 |

