## Supplementary figures and images for "Corrective sub-movements link feedback to feedforward control in the cerebellum"

### Supplemental figure 1

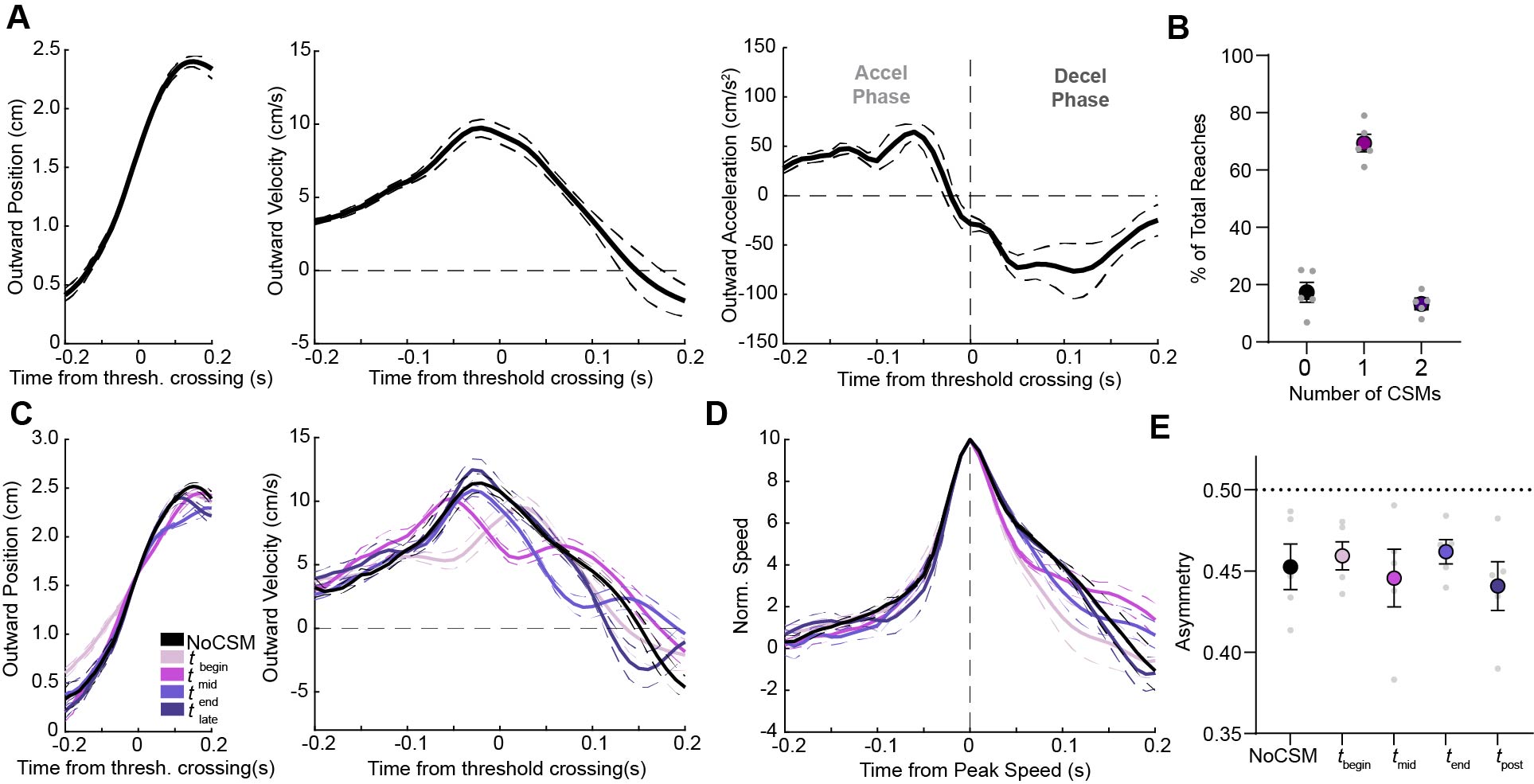

### Supplemental figure 2

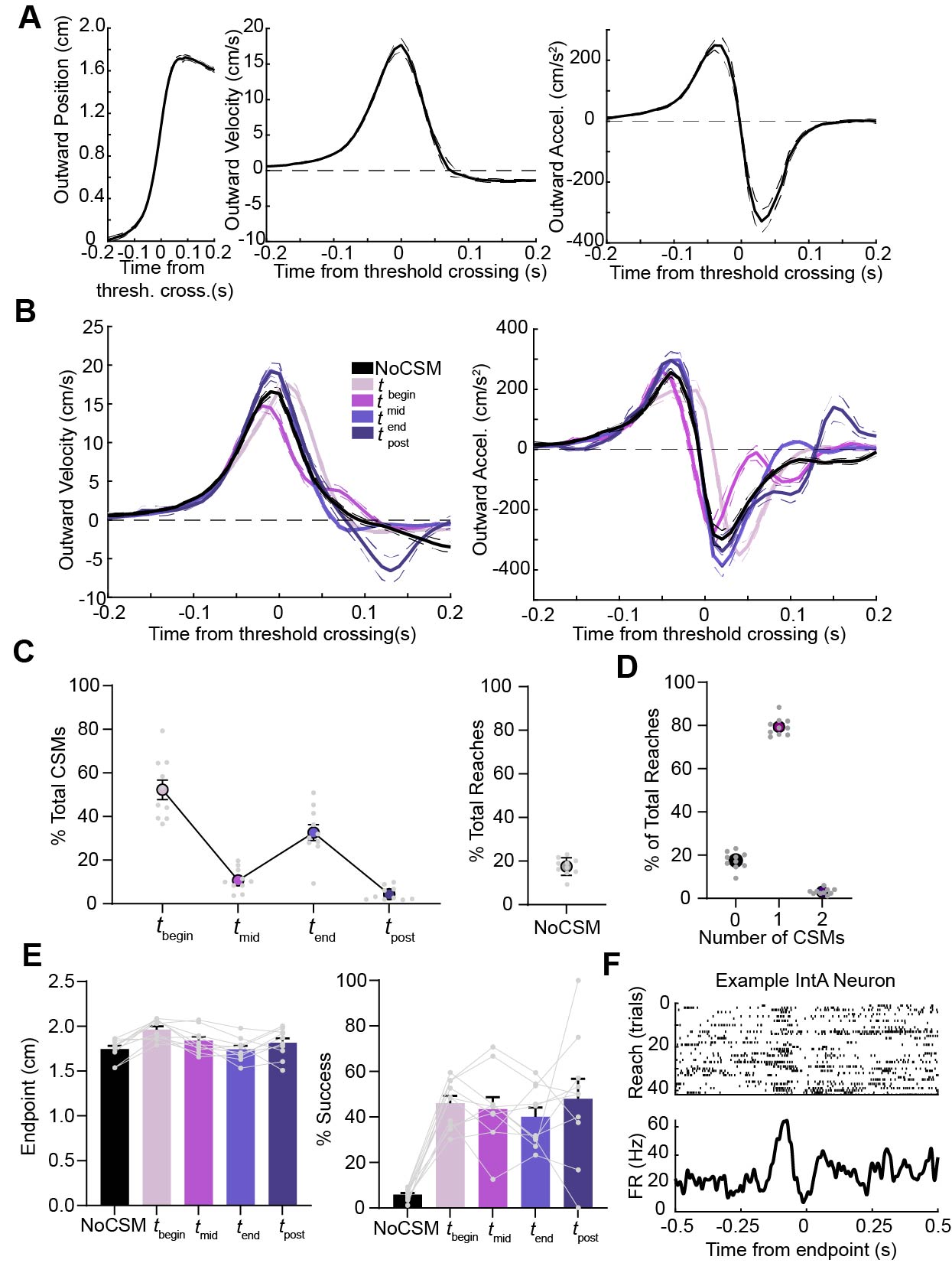

### Supplemental figure 3

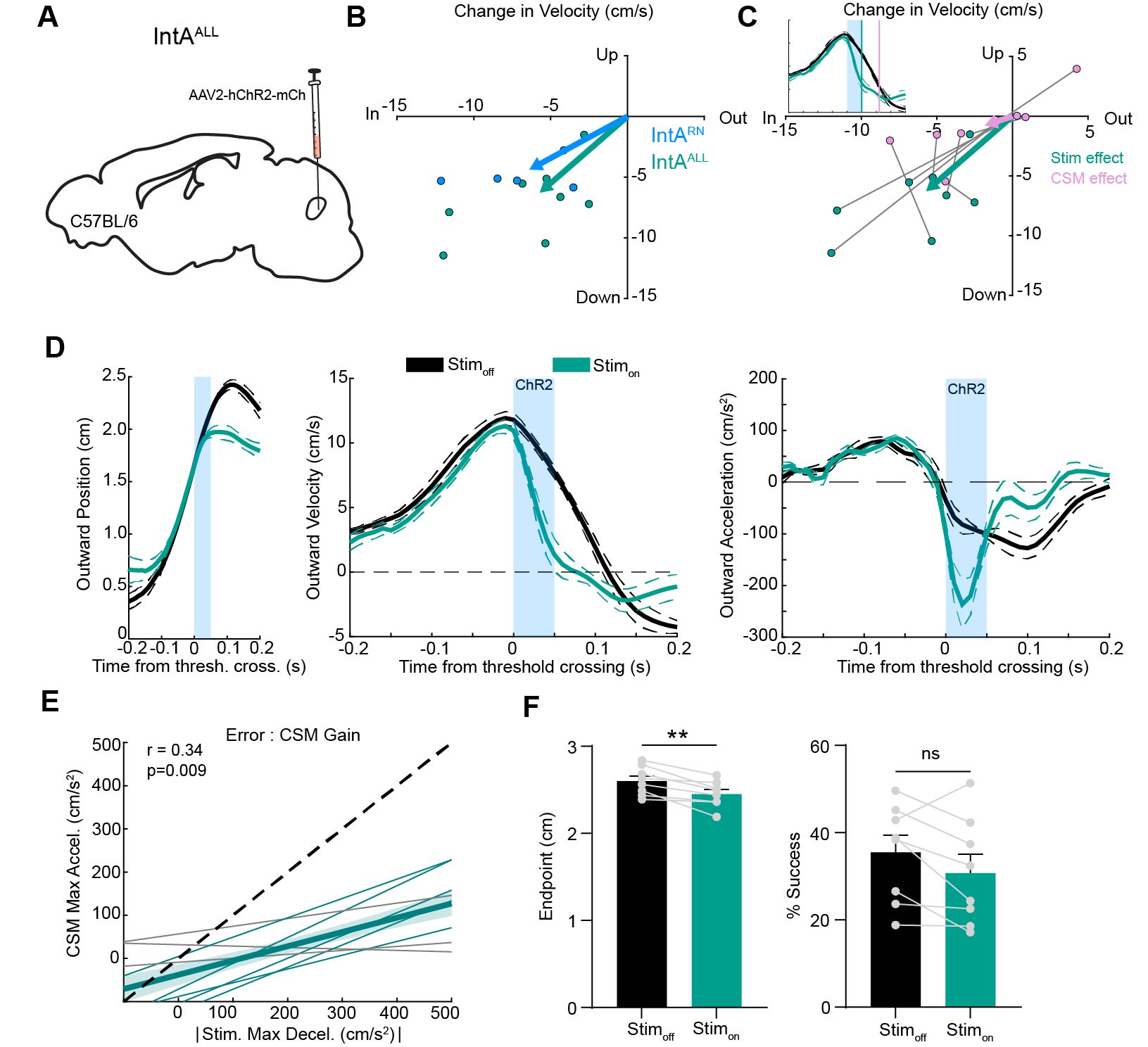

### Supplemental figure 4

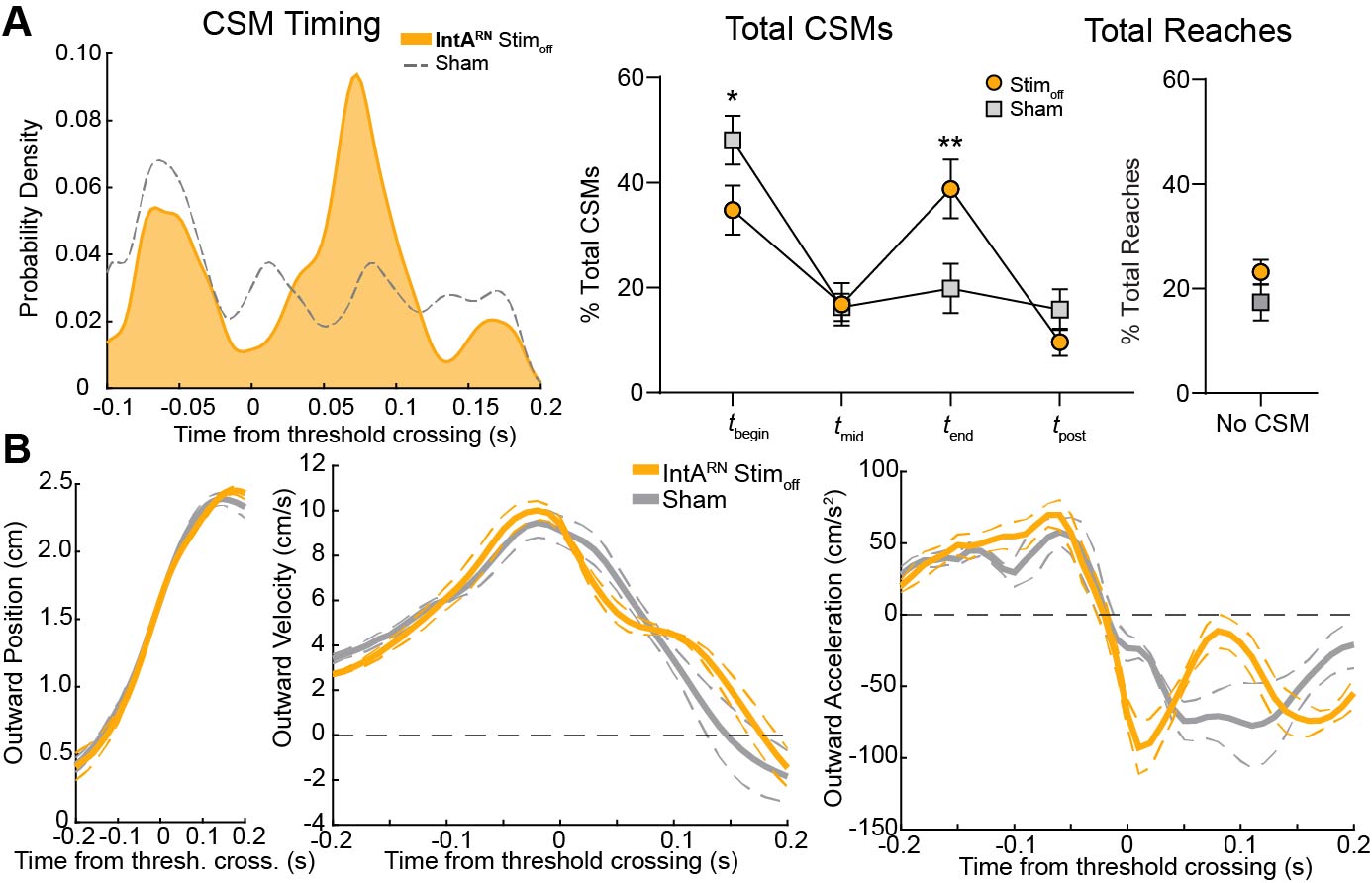

### Supplemental figure 5

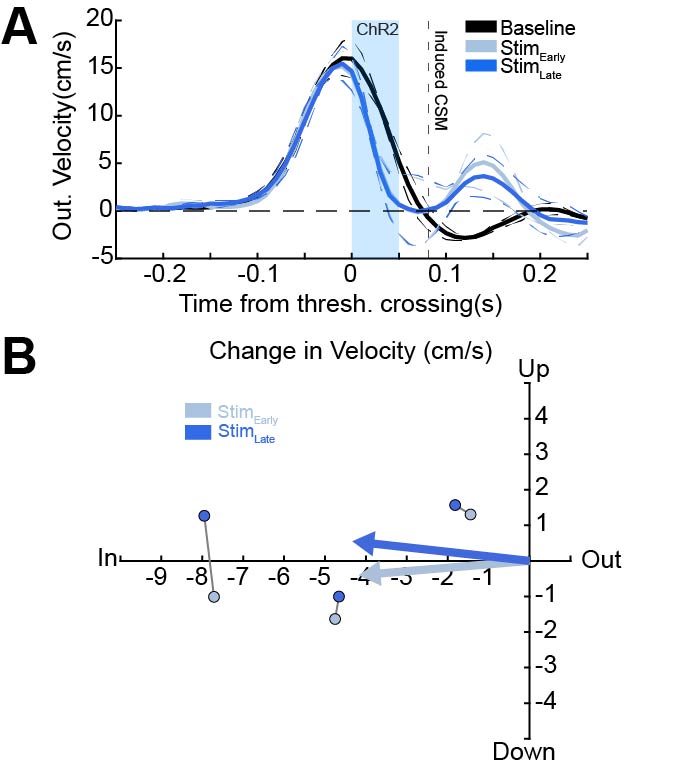
